# A role for neutral variation in the evolution of C_4_ photosynthesis

**DOI:** 10.1101/2020.05.19.104299

**Authors:** Shanta Karki, HsiangChun Lin, Florence R Danila, Basel Abu-Jamous, Rita Giuliani, David M Emms, Robert A Coe, Sarah Covshoff, Helen Woodfield, Efren Bagunu, Vivek Thakur, Samart Wanchana, Inez Slamet-Loedin, Asaph B. Cousins, Julian M Hibberd, Steven Kelly, W Paul Quick

**Author notes:** These authors contributed equally to this work (Shanta Karki, HsiangChun Lin). Authors for correspondence ***Corresponding Authors* Name:** Julian M. Hibberd **Email:** **Name:** Steven Kelly, **Email:**, **Name:** W. Paul Quick, **Email:**. ***Author contributions*** JMH and WPQ designed the study. BAJ, DME, SW, VT and SKe conducted the bioinformatic and phylogenetic analysis. SKa, ISL, HSL and WPQ produced and verified the transgenic lines. RC and HSL undertook physiological and metabolic analysis. RG and ABC conducted gas exchange analysis on transgenic plants. FD performed the confocal microscopy. SKe and WPQ analysed the data. SKe wrote the manuscript. WPQ, JMH and RAC edited the manuscript. All authors read and approved the final manuscript.

## Abstract

Convergent trait evolution is a recurrent phenomenon in all domains of the tree of life. While some convergent traits are caused by simple sequence changes, many are associated with extensive changes to the sequence and regulation of large cohorts of genes. It is unknown how organisms traverse this expansive genotype space to assemble such complex convergent phenotypes. C_4_ photosynthesis is a paradigm of large-scale phenotypic convergence. Conceptual and mathematical models propose that C_4_ photosynthesis evolved from ancestral C_3_ photosynthesis through sequential adaptive changes. These adaptive changes could have been rapidly assembled if modifications to the activity and abundance of enzymes of the C_4_ cycle was neutral in C_3_ plants. This neutrality would enable populations of C_3_ plants to maintain genotypes with expression levels of C_4_ enzymes analogous to those in C_4_ species and thus enable rapid assembly of a functional C_4_ cycle from naturally occurring genotypes given shared environmental selection. Here we show that there is substantial natural variation in expression of genes encoding C_4_ cycle enzymes between natural accessions of the C_3_ plant *Arabidopsis thaliana*. We further show through targeted transgenic experiments in the C_3_ crop *Oryza sativa*, that high expression of the majority of C_4_ cycle enzymes in rice is neutral with respect to growth, development, biomass and photosynthesis. Thus, substantial variation in the abundance and activity of C_4_ cycle enzymes is permissible within the limits of operation of C_3_ photosynthesis and the emergence of component parts of this complex convergent trait can be facilitated by neutral variation.

## Introduction

Common responses to environmental selection have produced a multitude of convergent complex traits in of all domains of the tree of life [1,2]. These complex traits include adaptations of existing anatomy such as the independent evolution of wings from flightless forelimbs in pterosaurs, birds and bats [3]. They also include the *de novo* formation of entire organs such as the evolution of camera-style eyes in cephalopods, vertebrates, arachnids and cnidarians [4,5]. For any such large complex traits, it has been challenging to determine the extent to which their emergence in independent lineages share a common genetic basis [1]. However, in recent times insight has been gained through study of genome sequences. Examples include the shared genome-wide signatures of convergent evolution in echolocating mammals [6], the molecular basis of coloration in vertebrates [7], and the changes in gene expression that underlie C_4_ photosynthesis [8–11]. While these studies have identified many of the molecular components that contribute to the complex convergent phenotypes, they do not provide insight into molecular mechanisms through which these phenotypes emerge. Moreover, the number of genes they identify highlights the combinatorial problem of how populations of organisms traverse the expansive genotype space required for the assembly of the complex convergent phenotype.

The C_4_ photosynthetic pathway is a paradigm for the evolution of complex convergent traits, requiring changes to organismal anatomy, development and physiology [12]. With at least 65 independent origins distributed across the angiosperms [13], it is ideally suited to investigate the mechanisms through which such convergent traits are assembled. The repeated evolution of the pathway is thought to have occurred as an adaptation to a drop in atmospheric CO_2_ concentration that occurred 35 million years ago [4]. The set of biochemical, physiological and anatomical changes that contribute to this adaptation counteract the reduction in atmospheric CO_2_ and also result in significant energy [14], water and nitrogen [15,16] savings [5]. As a result, C_4_ species tend to have increased productivity in tropical and sub-tropical habitats, and today C_4_ species represent some of the world’s most productive crops [17,18]. Concerns about future food security have led to the suggestion that C_4_ photosynthesis should be introduced into C_3_ crops such as rice to increase their yield potential [19,20]. This is an ambitious goal, and understanding the mechanisms by which the C_4_ pathway evolved in disparate C_3_ lineages holds potential to help guide these efforts.

Substantial insights into the evolution of C_4_ photosynthesis have been obtained through comparative analysis of genes, genomes, transcriptomes, ecological traits and physiological properties of C_3_ and C_4_ plants [21–24]. These studies have collectively revealed that all enzymes required to conduct C_4_ photosynthesis are present in C_3_ plants, however their abundance, activity, kinetic properties and subcellular localisation are altered in C_4_ species [22,25–27]. This set of biochemical changes occurs in concert with a substantial change in leaf anatomy [28], such that C_4_ plants have reduced vein spacing, altered cell-type specification around veins, and novel cell-specific organelle function when compared to C_3_ plants [28]. Thus, it is likely that multiple changes to both coding and regulatory sequences of multiple genes with developmental and biochemical functions are required to evolve a functional C_4_ cycle.

Despite this phenotypic complexity some annual plant lineages managed to evolve C_4_ photosynthesis relatively quickly (~5 million generations [29]), while others have been slower (~30 million generations [30]) or have not evolved C_4_ photosynthesis [22]. This recurrent evolution in a diverse array of plant groups has raised the question of how such a complex set of anatomical, biochemical and physiological changes can evolve on so many separate occasions. Conceptual, mathematical and statistical models of C_4_ evolution suggest that this suite of changes does not happen all at once. Instead, multi-step evolutionary trajectories likely occur such that successive adaptive changes traverse a phenotypic fitness path from C_3_ to C_4_ photosynthesis [31–34].

Evolutionary trajectories that bridge the gap between C_3_ and C_4_ photosynthesis could be rapidly traversed if variation in the activity and abundance of enzymes required to carry out the C_4_ cycle was neutral with respect to the native, physiological function in C_3_ plants. Such variation coupled to phenotypic neutrality is commonly referred to as “cryptic genetic variation” and is thought to be a major facilitator of trait evolution across all domains of life [35–37]. This is because genetically diverse populations are more likely to contain genotypes with a range of activity and abundance in phenotypically neutral enzymes. Such ‘pre-adapted’ phenotypes may confer an immediate selective advantage when the environment changes and a new selection pressure emerges [38–42]. Thus, in the context of C_4_ photosynthesis, neutrality (or near neutrality) of variance in the activity or abundance of enzymes of the core C_4_ cycle could enable rapid assembly of a functional C_4_ cycle from naturally occurring variation in a population. Moreover, such cryptic genetic variation could help explain the extent of the recurrent evolution of this complex convergent trait.

Here we test the hypothesis that variation in the activity and abundance of enzymes in the C_4_ pathway is neutral in a C_3_ background. Through analysis of natural variation in the C_3_ model plant *Arabidopsis thaliana* and single C_4_ cycle gene overexpression studies in the C_3_ crop plant *Oryza sativa* (rice), we show that none of the five enzymes required for a minimal NADP-ME C_4_ cycle adversely affects photosynthesis, growth or the global transcriptome. Thus, the enhanced activity and abundance of C_4_ cycle enzymes, in the correct cellular and sub-cellular context for C_4_ photosynthetic function, is neutral in this C_3_ context. These findings suggest that cryptic genetic variation may have facilitated the recurrent adaptive evolution of C_4_ photosynthesis.

## Results

### Genes encoding C_4_ cycle enzymes have high levels of variation in expression and molecular sequence evolution in C_3_ plants

The minimal set of enzymes required to conduct a typical NADP-ME C_4_ cycle, comprises carbonic anhydrase (CA), phospho*enol*pyruvate carboxylase (PEPC), malate dehydrogenase (MDH), NADP-malic enzyme (NADP-ME) and pyruvate, orthophosphate dikinase (PPDK) (Figure 1). To assess whether the genes encoding these enzymes exhibit high variance in expression within populations of C_3_ plants, a transcriptomic analysis of 17 different natural accessions of *Arabidopsis thaliana* [43] was interrogated. Equivalent datasets do not exist for other species. Variance in relative mRNA abundance among natural accessions was assessed and the variance percentile of the core C_4_ genes, with respect to all expressed genes, was evaluated. *Arabidopsis thaliana* orthologs of the core C_4_ genes were identified using a phylogenetic approach (see methods). In the case where multiple paralogs of a core C_4_ gene are present in the genome of *Arabidopsis thaliana*, then the paralog that was recruited to function most frequently in NADP-ME C_4_ eudicot species [10] is shown and data for all paralogs are provided in Supplemental File S1. In all cases there was substantial variation in the abundance of transcripts encoding the core C_4_ cycle enzymes (Figure 2A, Supplemental File S1). The least variable gene, *AtPPDK*, was more variable than 84% of expressed genes in the *Arabidopsis* leaf, while the most variable, *AtCA*, was in the 98^th^ percentile. Thus, the abundance of transcripts encoding core enzymes of the C_4_ cycle are highly variable between natural accessions, suggesting that variation in abundance of these enzymes is phenotypically neutral under the growth conditions used here.

**Figure 1.**
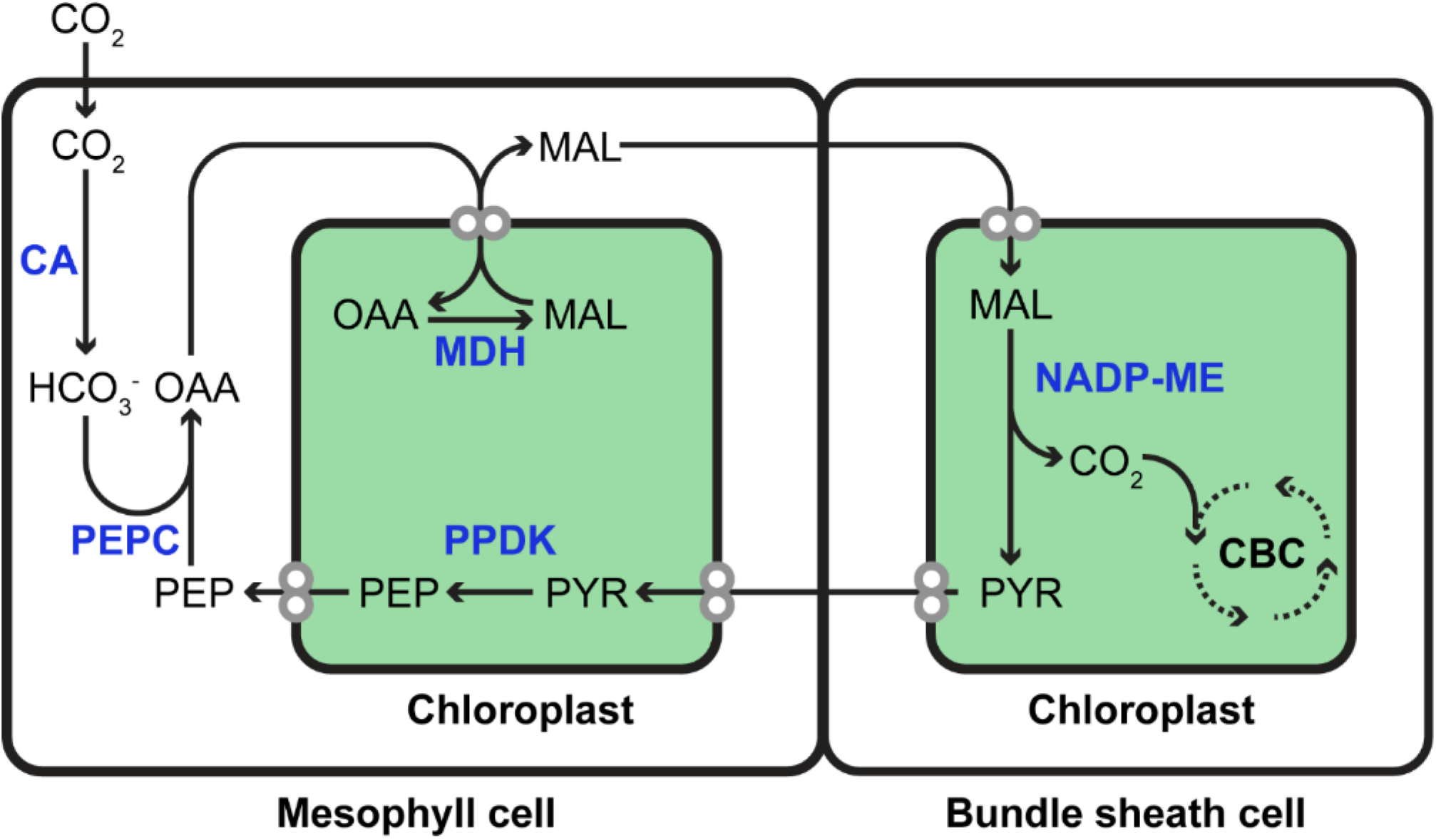
Cartoon of the enzymes required to carry out a minimal NADP-ME C_4_ cycle. Enzymes are depicted in bold blue font, metabolites in black font. CA, carbonic anhydrase. PEPC, phospho*enol*pyruvate carboxylase. MDH, malate dehydrogenase. NADP-ME, NADP-malic enzyme. PPDK, pyruvate, orthophosphate dikinase. OAA, oxaloacetate. MAL, malate. PYR, pyruvate. PEP, phospho*enol*pyruvate.

**Figure 2.**
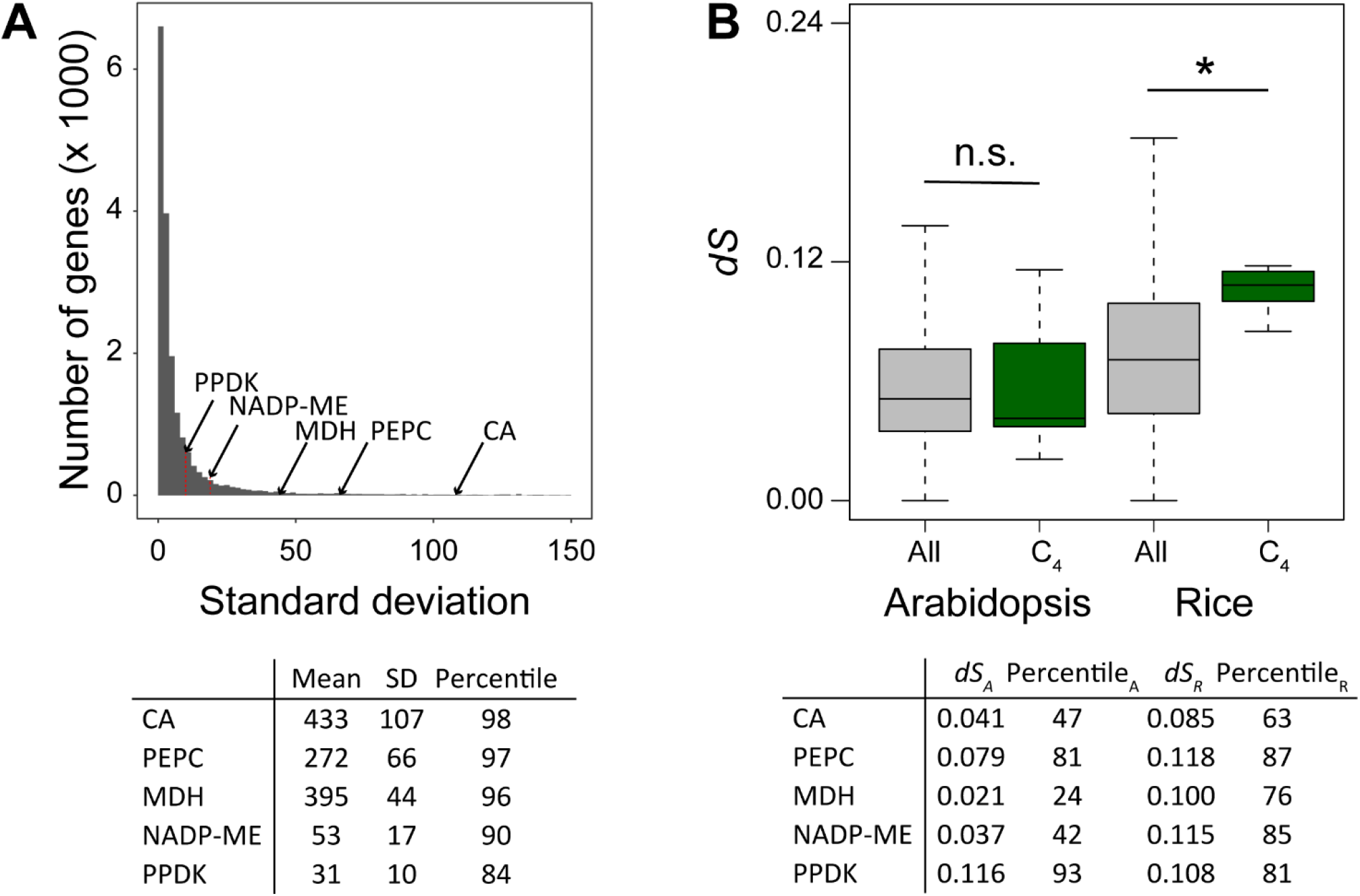
Analysis of variance in relative mRNA abundance and molecular sequence evolution of genes encoding core C_4_ cycle enzymes in *Arabidopsis thaliana* and *Oryza sativa*. **A)** Histogram of standard deviation of all expressed genes for 17 different natural accessions of *Arabidopsis thaliana*. Position of core C_4_ cycle genes is indicated by arrows. For each of the core C_4_ cycle genes the mean and standard deviation and percentile rank is provided in a table below the plot. The mean variance of this cohort is significantly larger than would be expected by chance (p < 0.001, Monte Carlo resampling test). The complete expression dataset is also provided as Supplemental File S1. **B)** Box plots of *dS* (number of synonymous substitutions per synonymous site per gene that occur in the whole natural or cultivated accession dataset) for all genes in the genome as compared to the cohort of core C_4_ cycle genes. For each of the core C_4_ cycle genes the *dS* value and its percentile rank are given. * indicates significant difference (t-test, p < 0.01).

Given that transcripts encoding core enzymes of the C_4_ cycle exhibited significantly higher variance in abundance between natural accession of *Arabidopsis thaliana* than would be expected by chance, we hypothesised that the genes encoding these enzymes may also be subject to enhanced rates of neutral variation. To test this hypothesis, we analysed the number of within-species synonymous substitutions in the coding sequences of these genes in the 1001 Arabidopsis genome project [44]. Although natural accession data of this magnitude was not available for other species at the time of this analysis, a similar scale dataset was available for cultivated accessions of *Oryza sativa* [45]. This revealed that while *AtPEPC* and *AtPPDK* exhibited high levels of neutral variation, on average the cohort of genes in natural accessions of *Arabidopsis thaliana* had levels of neutral variation that are representative of the genome-wide distribution (Figure 2B). In contrast, levels of neutral variation for the cohort of core C_4_ cycle genes in rice were significantly higher than expected (Figure 2B).

### An experimental system for testing the neutrality of C_4_ cycle enzymes in a C_3_ context

Given that the above analysis suggested that variation in core C_4_ cycle genes was phenotypically neutral in a C_3_ context, we sought to directly test this hypothesis using a transgenic approach. These transgenic tests were conducted in the model C_3_ species *Oryza sativa spp. Indica* cultivar IR64 [46], as it is of direct relevance to C_4_ engineering efforts [19,20]. Independent transgenic lines were generated to overexpress each of the six core C_4_ cycle enzymes. For each enzyme, a genomic clone comprising the complete maize gene sequence, including the promoter region and all introns and exons, was chosen for overexpression (Table 1). This strategy was selected for two reasons. 1) The biochemical function of each of the maize enzymes has been experimentally validated, whereas not all of the enzymes encoded by the orthologous genes in rice have been characterised [47]. 2) Sequence differences between the maize and rice genes enables transcripts attributable to the endogenous and heterologous genes to be distinguished so that relative abundances can be measured. The exception to this rule was the construct for overexpression of carbonic anhydrase (Table 1). Here, due to complexity in developing a specific antibody for maize cytosolic β-Carbonic Anhydrase (CA), the coding sequence (CDS) was translationally fused to a C-terminal AcV5 epitope tag and expressed under the control of the maize PEPC promoter (ZmPEPC_pro_). In all cases, three independent single insertion transgenic lines with the highest transgene expression (assessed by both transcript and protein quantification) were taken forward for subsequent analysis (Supplemental Figure S1 and S2). In the case of malic enzyme, protein expression was only detected in a single transgenic line which contained ≥6 copies of the transgene construct. Thus, this was the only malic enzyme line taken forward for further analysis.

**Table 1:** Construct details.

|  | Accession | CDS or genomic | Promoter |
| --- | --- | --- | --- |
| CA | GRMZM2G348512 | CDS | ZmPEPC |
| PEPC | GRMZM2G083841 | genomic | ZmPEPC |
| MDH | GRMZM2G129513 | genomic | ZmMDH |
| NADP-ME | GRMZM2G085019 | genomic | ZmME |
| PPDK | GRMZM2G306345 | genomic | ZmPPDK |

### Overexpressed C_4_ cycle enzymes localise to the correct cellular compartment context in transgenic rice plants

Given that protein of the correct size was expressed in each of the transgenic lines, we subsequently sought to determine whether those proteins were localised to the correct cell type and subcellular compartment. It was not possible to conduct immunolocalisation analysis using the anti-MDH and anti-ME antibodies as these antibodies cross-reacted with native protein in wild-type rice and so the endogenous and exogenous proteins could not be distinguished from each other (Supplemental Figure S3). However, for the other transgenes immunolocalisations revealed that each overexpressed protein accumulated preferentially in the same cell type and subcellular compartment as the endogenous gene in *Zea mays* (Figure 3). Specifically, ZmCA2 (Figure 3A and 3B) and ZmPEPC (Figure 3C and 3D) accumulated in the cytosol of mesophyll cells, while ZmPPDK localised to chloroplasts in both bundle sheath and mesophyll cells (Figure 3E and 3F). Although PPDK activity is only required in chloroplasts of the mesophyll, in maize transcripts encoding PPDK and also the enzyme itself accumulate to significant levels in mesophyll and bundle sheath cells [48–50]. This observed localisation, while consistent with the expectation from maize, is in contrast to previous studies that have reported that the *ZmPPDK* promoter drives GUS gene expression in mesophyll cells, with little expression in bundle sheath cells (Matsuoka et al., 1993). Thus, it appears that there are differences between transcript accumulation driven by the promoter alone and transcript/protein accumulation driven by the intact gene. Overall, localisation of proteins in the transgenic rice lines were consistent with the patterns of gene expression and protein accumulation in C_4_ maize.

**Figure 3.**
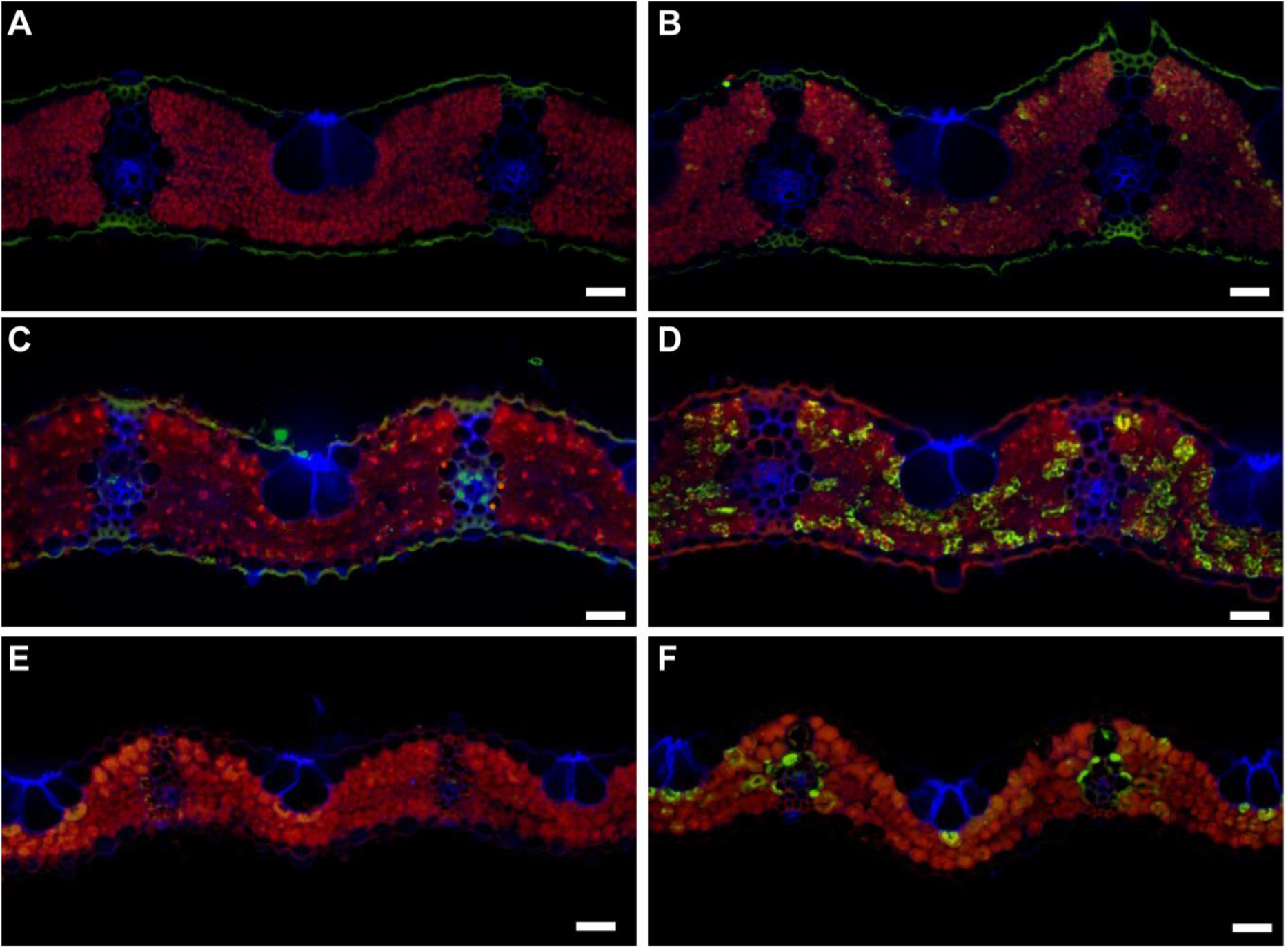
Localisation of overexpressed proteins. A & B) Rice leaf cross sections stained with anti-AcV5 antibody for A) an un-transformed wild-type plant and B) a transgenic line overexpressing ZmCA2-AcV5. C & D) Rice leaf cross sections stained with anti-ZmPEPC antibody for B) an un-transformed wild-type plant and B) a transgenic line overexpression ZmPEPC. E & F) Rice leaf cross sections stained with anti-ZmPPDK antibody for E) an un-transformed wild type plant and F) a transgenic line overexpressing ZmPPDK. Magnification: 200x. Scale bar: 20 μm.

### Overexpressed enzymes confer enhanced enzyme activity

Given that the overexpressed transgenes resulted in protein that localised to the anticipated cell types and subcellular compartments, we next investigated whether this also resulted in additional enzyme activity within the leaf. In the case of carbonic anhydrase, the overexpression of ZmCA2 did not result in significantly elevated levels of carbonic anhydrase activity per unit leaf area (Figure 4A). This lack of an observed increase was likely due to the high levels of carbonic anhydrase activity that are present in wild type rice leaves [51]. In contrast, PEPC activity in rice leaves is low and overexpression of *ZmPEPC* resulted in a significant increase in leaf level PEPC activity (Figure 4B). Similarly, significantly elevated levels of MDH (Figure 4C) and ME (Figure 4D) activity per unit leaf area were also detected in the respective transgenic lines. Although, expression of PPDK enzyme was detected in all transgenic lines, enhanced PPDK activity was only measured in two of the three independent single insertion transgenic lines overexpressing *ZmPPDK*. Thus, although ectopic accumulation of each protein could be readily detected, higher activities were only detected for four of the five C_4_ cycle enzymes analysed.

**Figure 4.**
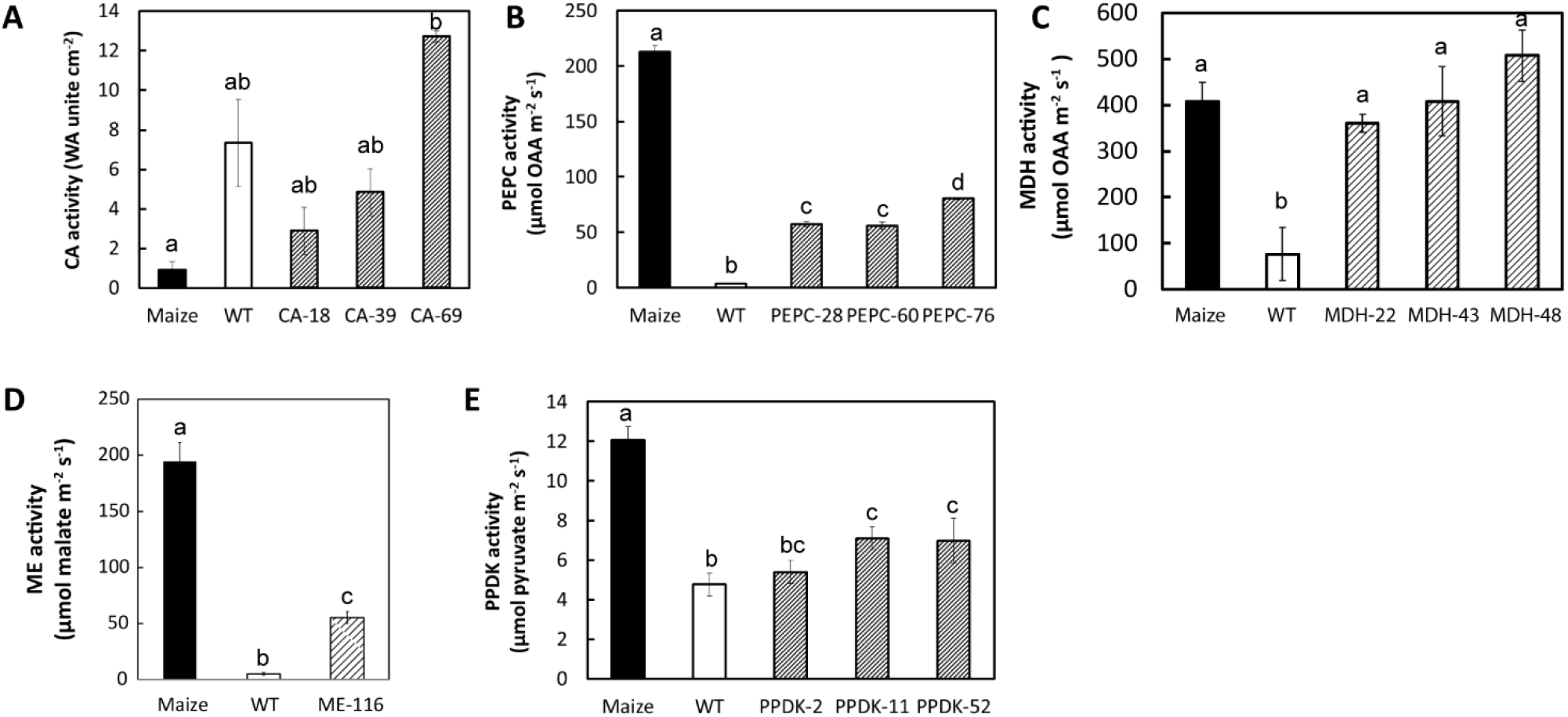
Leaf level enzyme activity assays transgenic lines overexpressing ZmCA2, ZmPEPC, ZmMDH, ZmME and ZmPPDK. In all cases enzyme assays from leaves of non-transformed wild-type rice plants (WT) and leaves of non-transformed wild-type maize plants (Maize) are shown for reference. A) Carbonic anhydrase. B) Phosphoenolpyruvatecarboxylase. C) Malate dehydrogenase. D) Malic enzyme. E) Pyruvate, orthophosphate dikinase. Different letters above bars indicate those values that are statistically different based on a one-way ANOVA with a Tukey multiple comparison test for post-hoc pairwise comparison (p < 0.05).

To assess whether these altered enzyme activity levels were similar to those observed for analogous enzymes functioning in a C_4_ cycle, enzyme assays were also conducted on leaves of maize. This revealed that levels of MDH activity in leaves of the transgenic lines were analogous to those observed in maize leaves (Figure 4C). In contrast, levels of ME (Figure 4D) and PPDK (Figure 4E) were ~50% of what are observed in maize, and levels of PEPC were 25% of those found in maize. Thus, enzyme activity levels for the majority of the enzymes were elevated relative to wild-type rice leaves and are comparable to the levels observed in maize leaves.

### No phenotypic perturbation associated with overexpression of C_4_ cycle genes in Oryza sativa

Given that the enzymes were expressed, active and localised to the correct cell-type and subcellular localisation it was next investigated the extent to which this activity was detrimental to growth or photosynthesis in the transgenic plants. None of the transgenic lines showed altered chlorophyll content (Figure 5A), tiller number (Figure 5B), plant height (Figure 5C) or dry biomass (Figure 5D). Moreover, all plants were phenotypically indistinguishable compared with wild type controls (Supplemental Figure S4). The one exception to this rule was the transgenic line overexpressing *ZmME*. This line exhibited a small decrease in plant height (Figure 5C), however, this difference did not result in a detectable difference in dry biomass (Figure 5D). In cases where an individual line differed from wild-type or from other independent transgenic lines, this difference did not correlate with differences in enzyme activity (Figure 4) and was thus likely attributable to somaclonal variation during callus regeneration. Overall, transgene expression caused no perturbation to growth or biomass.

**Figure 5.**
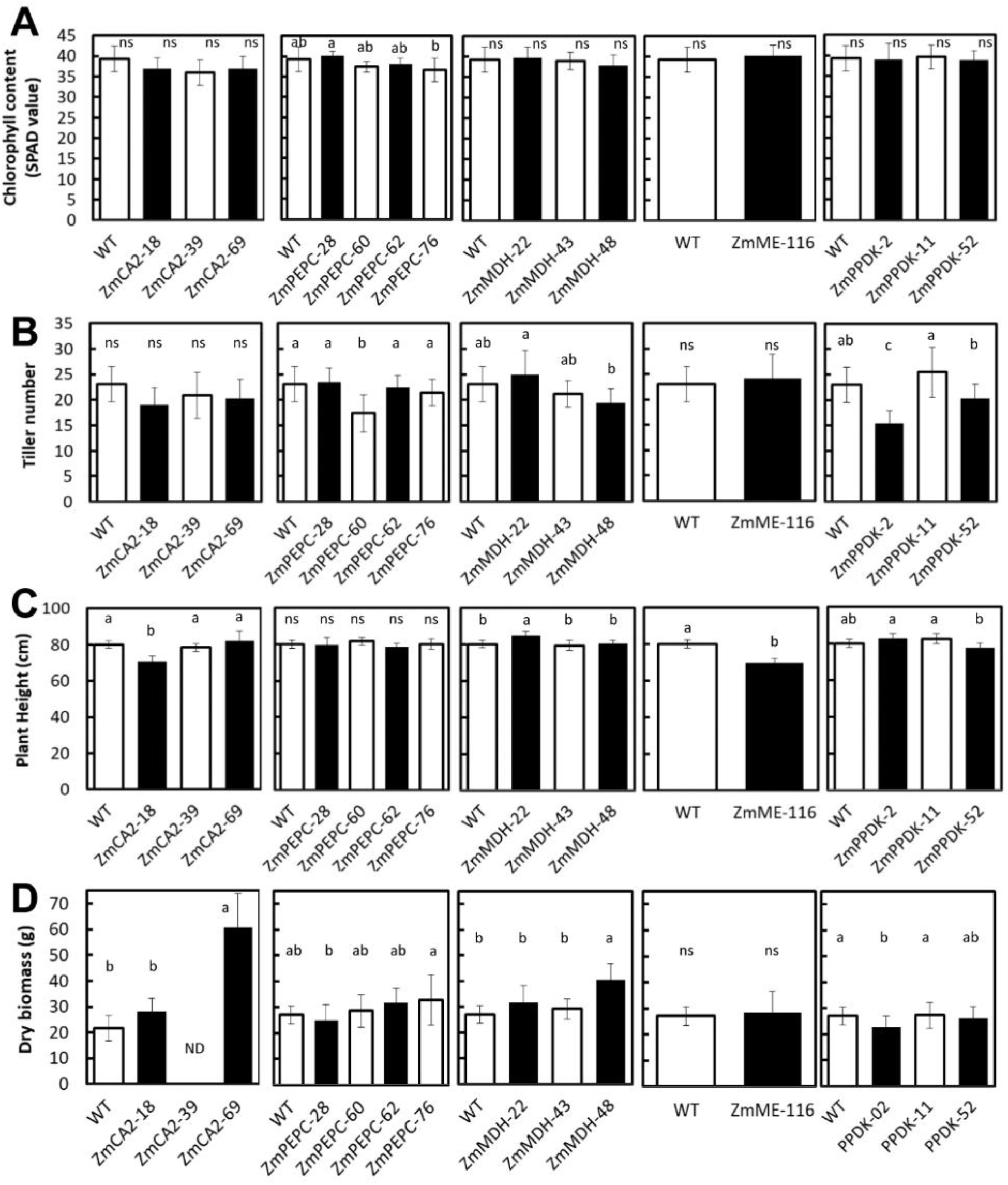
Leaf Chlorophyll content, tiller number, plant height and dry biomass of wild-type and transgenic lines overexpressing ZmCA2, ZmPEPC, ZmMDH, ZmME and ZmPPDK grown in rice paddy fields at IRRI. Chlorophyll SPAD values are the average ± SE of three leaves from 15 plants using the upper youngest fully expanded leaves at maximum tillering stage; 70 days post-germination. Tiller number and plant height are the average ± SE of 15 individual plants measured at maximum tillering stage; 70 days post-germination. Dry biomass is the total dry weight of leaf, stem, sheath tissues and panicles at harvesting stage; 98 days post-germination. Dry biomass is the average ± SE of 20 individual plants, except 10 plants for ZmCA2 lines. ND indicates that dry biomass was not determined in ZmCA2-39 line. Different letters above bars indicate those values that are statistically different based on a one-way ANOVA with a Tukey multiple comparison test for post-hoc pairwise comparison (p-value<0.05). “ns” indicates non-significant.

For PEPC and PPDK the field studies presented here are consistent with controlled environment experiments which showed that overexpression of *ZmPEPC* or *ZmPPDK* in rice did not affect plant growth [52]. However, it has previously been reported that expression of *ZmME* in rice enhanced photoinhibition leading to photo-bleaching and stunted growth in green house conditions [53]. It is possible, that differences in growth conditions (field vs greenhouse), or differences in enzyme abundance, or differences in the cell type specificity of the promoters used (i.e. we used the mesophyll cell specific PEPC promoter whereas previous studies used the ubiquitous CaMV 35S or cab promoter) are responsible for this difference in phenotype. Regardless, the data presented here show that rice plants with significantly enhanced ME activity (i.e. ~45% of that found in maize and 10 fold higher than wild type rice) had no aberrant growth phenotype under field conditions.

### No photosynthetic perturbation associated with overexpression of C_4_ cycle genes in Oryza sativa

The lack of perturbation to growth in field conditions described above is consistent with the hypothesis that variation in the abundance and activity of enzymes of the C_4_ cycle is neutral in a C_3_ context. However, these field experiments were conducted under current atmospheric conditions (~420ppm CO_2_). To provide insight into whether such neutrality would also be manifest at atmospheric conditions that would have been present prior to the emergence of C_4_ photosynthesis (i.e. ≥1000ppm CO_2_) or during the emergence of C_4_ photosynthesis (i.e. ~200ppm CO_2_), the response of photosynthesis to altered CO_2_ levels was measured. In each case, there was no significant difference in light-saturated photosynthetic rate at sub-ambient CO_2_ concentrations typical of the period during which the ≥60 independent C_4_ lineages evolved (Figure 6). Furthermore, for transgenic lines overexpressing *ZmCA* (Figure 6A), *ZmPEPC* (Figure 6B) and *ZmMDH* (Figure 6C) there was no significant difference in light-saturated photosynthetic rates at elevated CO_2_ concentrations typical of the period prior to the emergence of C_4_ photosynthesis. Additional tests were conducted on two of lines overexpressing *ZmCA* using online carbon and oxygen isotope discrimination analysis [54,55] to confirm that cytosolic CA activity did not affect other photosynthetic parameters such as mesophyll conductance (Supplemental File S2). In contrast, there appeared to be some reduction in maximal photosynthetic rate at elevated CO_2_ concentrations in transgenic lines overexpressing *ZmME* (Figure 6D) or *ZmPPDK* (Figure 6E). Thus, it is possible that variation in the abundance and activity of these specific C_4_ enzymes may have detrimental effects on growth at elevated CO_2_ concentrations. However, for the majority of the enzymes of the C_4_ cycle, variation in the abundance and activity of these enzymes had no effect on growth, and no effect on photosynthesis under current atmospheric conditions or under conditions typical of the periods both prior to and during the emergence of C_4_ photosynthesis. This result further supports the hypothesis that variation in the abundance and activity of enzymes of the C_4_ cycle is relatively neutral in a C_3_ context.

**Figure 6.**
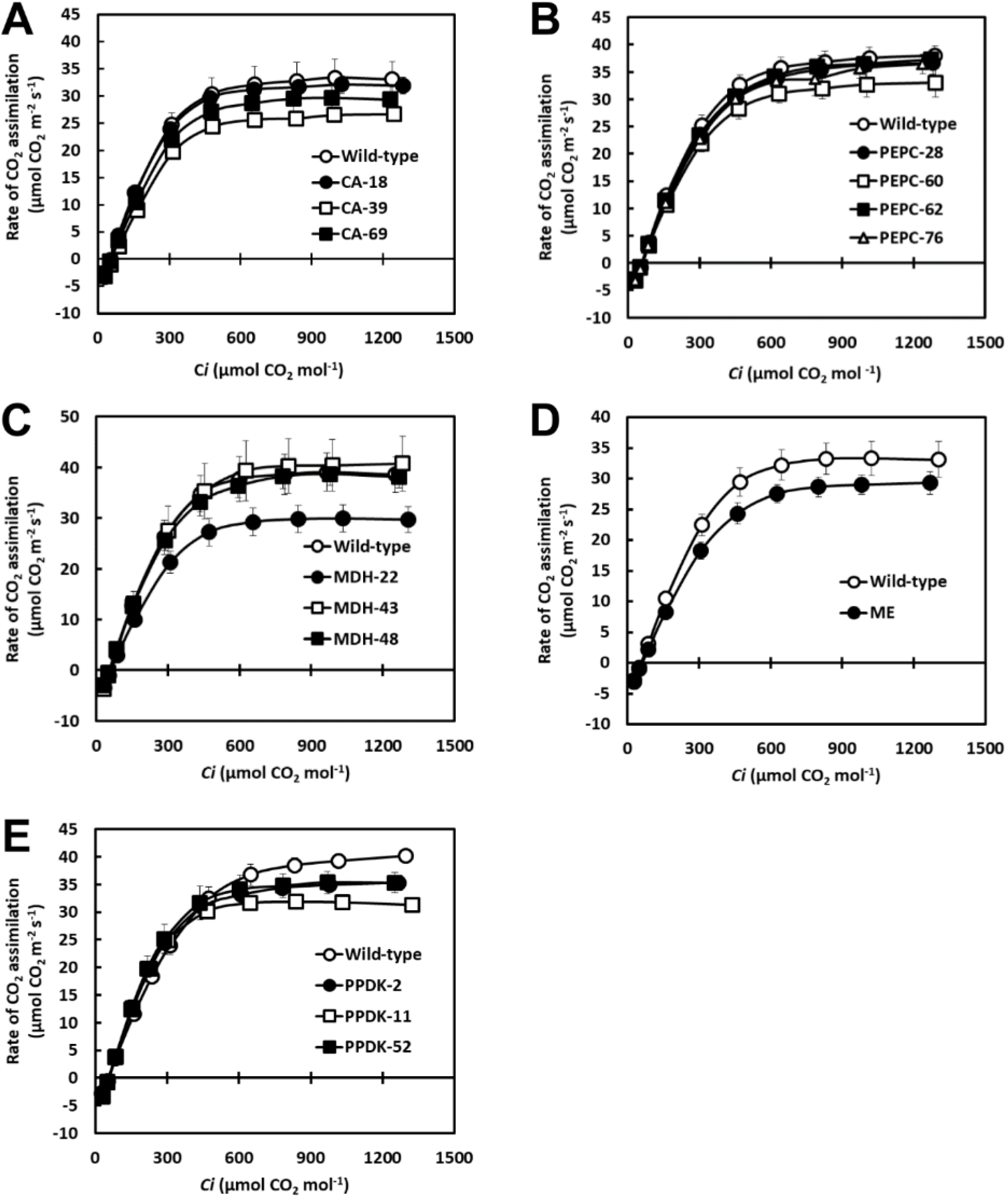
CO_2_ response curves for rice transgenic lines overexpressing enzymes of the C_4_ cycle. In all cases *C_i_* is the sub-stomatal CO_2_ concentration within the leaf. A wild-type non transformed control is included in each plot. A) Transgenic lines overexpressing ZmCA. B) Transgenic lines overexpressing ZmPEPC. C) Transgenic lines overexpressing ZmMDH. D) Transgenic lines overexpressing ZmME. E) Transgenic lines overexpressing ZmPPDK.

### The absence of growth and photosynthetic phenotypes is not attributable to compensatory mechanisms operating at a transcriptome-wide level

Given the lack of perturbation to growth or photosynthesis in these transgenic lines we sought to determine whether this apparent neutrality was attributable to compensatory mechanisms operating at the level of gene expression. Specifically, could alteration of the expression of genes encoding enzymes or transporters in related biochemical pathways (or which catalyse the reverse reactions) help explain the lack of a detrimental effect? To test this, each of the transgenic lines were subject to transcriptome sequencing and differential expression analysis.

As expected from the protein expression and enzyme assay data, transcripts corresponding to the introduced transgenes were detected in each of the transgenic lines (Figure 76A). In all cases, overexpression of the maize gene (Figure 7A, boxes labelled X) did not cause compensatory change in the abundance of the endogenous ortholog (Figure 7A, boxes labelled E), as these values were not significantly different to those in wild type plants (Figure 7A, boxes labelled W). With the exception of the transgenic lines overexpressing *ZmCA*, the transcript abundance of the overexpressed maize genes (Figure 7A, boxes labelled X) were higher than the transcript abundance of the endogenous gene (Figure 7A, boxes labelled E). For the transgenic lines overexpressing *ZmCA, ZmPEPC* and *ZmMDH*, fewer than 22 genes were detected as differentially expressed when transcript profiles were compared to wild type plants (Figure 7B, Supplemental File S3). Of these, none encoded components of related biochemical pathways or proteins from which any functional significance could be inferred. When the transgenic lines overexpressing *ZmME* and *ZmPPDK* were compared to wild type plants there were 277 and 27 differentially expressed genes, respectively (Figure 7B, Supplemental File S3). Functional term enrichment analysis revealed an overrepresentation of light reaction and Calvin cycle genes in the sets of genes that were downregulated in the transgenic lines overexpressing *ZmME* and *ZmPPDK* (Supplemental File S3). Thus, there is some perturbation to the leaf transcriptome that is consistent with the detectable reduction in maximal photosynthetic rate at elevated CO_2_ concentrations in transgenic lines overexpressing *ZmME* or *ZmPPDK*. Overall, however, increased activity of core C_4_ cycle enzymes through overexpression appears to be within the permissible limits of variation for normal C_3_ photosynthesis.

**Figure 7.**
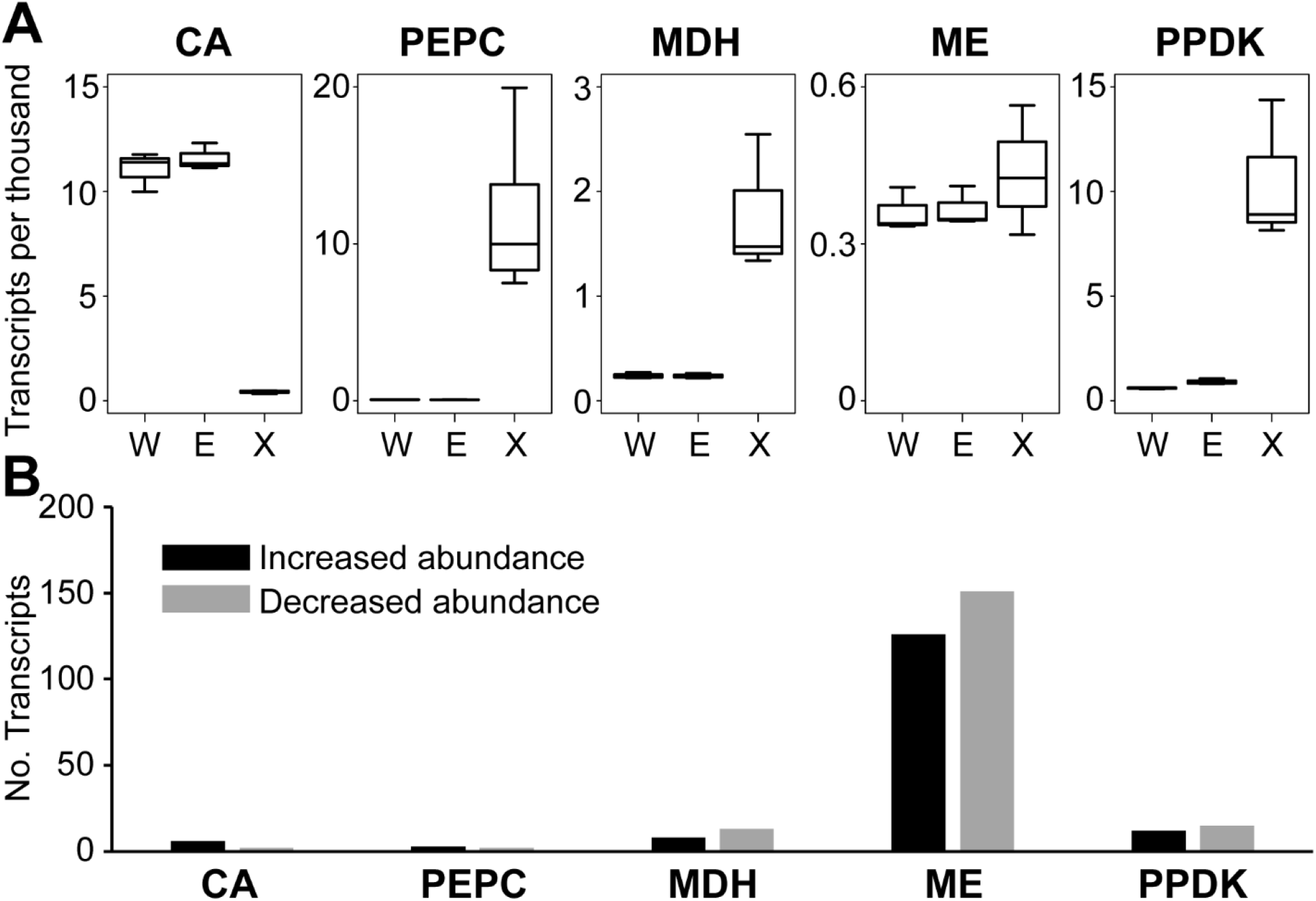
Transcriptome analysis of rice transgenic lines overexpressing enzymes of the C_4_ cycle. A) Comparison of the relative mRNA abundance of transcripts encoding C_4_ cycle enzymes. W is the transcript abundance of the endogenous rice gene in non-transformed rice plants, is the transcript abundance of the endogenous rice gene in the transgenic rice plants overexpressing the C_4_ cycle enzyme, and X is the transcript abundance of the exogenous maize gene encoding the C_4_ cycle enzyme. Data from transgenic lines overexpressing each enzyme is shown separately. Boxplots show the complete data range. The over expressed C_4_ cycle enzyme is shown above each plot. B) The number of upregulated and downregulated genes in the transgenic lines overexpressing enzymes of the C_4_ cycle. The y axis depicts the number of differentially express transcripts in that transgenic line in comparison to wild type rice plants.

## Discussion

The evolution of C_4_ photosynthesis is widely considered to be one of the most remarkable examples of phenotypic convergence in eukaryotic biology. In this work, we propose that cryptic genetic variation may have facilitated the rapid evolution of C_4_ photosynthesis. In support of this hypothesis we show that there is substantial natural diversity in the abundance of transcripts encoding C_4_ enzymes in the C_3_ plant *Arabidopsis thaliana,* and the majority of the biochemical differences that distinguish C_3_ and C_4_ plants can be recapitulated in C_3_ plants with limited or no effects on plant growth or photosynthesis. This finding means that the majority of the changes in abundance of enzymes required to conduct C_4_ photosynthesis can evolve independently without perturbation to growth within a population of C_3_ plants, and that these ‘pre-adapted’ phenotypes only require the correct series of crosses to assemble the complete set of enhanced enzyme abundances within one individual plant.

Although the change in the majority of the enzymes produced no detectable effect on growth or photosynthesis, a perturbation to photosynthesis was observed at elevated CO_2_ concentrations in lines overexpressing *ZmME* and *ZmPPDK*. Such a perturbation could mean that enhanced activity of these enzymes may have been detrimental under the high CO_2_ conditions prevalent prior to the emergence of C_4_ photosynthesis. If this was the case, it might be expected that standing genetic variation in the abundance of these enzymes would be reduced when compared to the other C_4_ cycle enzymes in wild populations of C_3_ plants. Consistent with this hypothesis we observed that variation in the transcriptome abundance of the genes encoding ME and PPDK is reduced when compared to the genes encoding other C_4_ cycle enzymes in different natural accessions of *Arabidopsis thaliana*. Thus, it is possible that acquisition of enhanced activity of these enzymes may be a limiting factor in the assembly of a C_4_ cycle from extant natural diversity.

Although, overexpression of PPDK exhibited a minor perturbation to photosynthesis at elevated CO_2_ this effect was not observed at ambient CO_2_ or at CO_2_ levels prevalent when C_4_ photosynthesis evolved. In this context, it is worth nothing that overexpression of PPDK during leaf senescence in either Arabidopsis or tobacco can be beneficial and lead to enhanced growth [56]. Such benefits to growth may help explain why modelling studies indicate that upregulation of PPDK may occur prior to upregulation of other C_4_ cycle enzymes [32].

The findings presented here do not negate or contradict other hypotheses concerning the timing or order of the evolution of C_4_ photosynthesis [31–34]. For example, the neutrality of C_4_ cycle enzyme abundance in a C_3_ context helps explain why the order of C_4_ trait evolution is flexible [32] and why these changes occur on a smooth fitness landscape [31]. It also provides new insight into the potential mechanisms by which these changes have occurred. Specifically, the findings presented here reveal that the acquisition of the biochemical functions required for the establishment of a C_4_ photosynthetic cycle do not need to evolve in series, and can instead evolve in parallel within a population and be assembled in series through sexual reproduction. The ability of such cryptic genetic variation to assist in the evolution of C_4_ photosynthesis may thus help to explain the high frequency of this phenotypic convergence.

## Methods

### Plant growth

Plants were grown in rice paddy fields at the International Rice Research Institute (IRRI), Los Baños Philippines, 14° 10019.900N, 121° 15022.300E. Seeds were placed at 50° C for 3 days and were then germinated in distilled water for 2 days. The germinated seeds were sown in seedling trays containing sterilized soil (taken from the IRRI upland farm) for 2 weeks and then transplanted into rice paddy fields. For gas exchange measurements, plants were grown in a screenhouse with a day/night temperature of 35/28 ± 3 °C.

### Generation of transgenic plants

To express high levels of ZmPEPC (GRMZM2G083841), ZmPPDK (GRMZM2G306345), ZmMDH (GRMZM2G129513) and ZmME (GRMZM2G085019) in rice, full-length genomic fragments encompassing the genes encoding these maize enzymes and their promoters were cloned into pSC0 vector (GenBank, Accession no. KT365905 [57]). Generation of pSC0/ZmPEPC vector was previously described [58]. A pSC0/ZmPPDK vector containing a full-length genomic fragment was created by subcloning ZmPPDK from pIG121Hm/PPDK [59] (a gift from Mitsue Miyao, NIAS, Japan) into pSC0. Gibson assembly was used to insert the gene into pSC0 vector. The necessary amplicons from the pIG121Hm/PPDK and pSC0 templates were amplified using Primer I: 5’-ATGCTCAACACATGAGCGAAGGGCCCATGACCATGATTACGCCAAG, Primer II: 5’-TGTGCATGTCGCTAGGATCCGGTACCGAATGCTAGAGCAGCTTGA, Primer III: TCAAGCTGCTCTAGCATTCGGTACCGGATCCTAGCGACATGCACA, Primer IV: 5’-CTTGGCGTAATCATGGTCATGGGCCCTTCGCTCATGTGTTGAGCAT. The full-length genomic sequences of ZmME and ZmMDH were amplified from BACs sourced from BACPAC resources (Children’s Hospital Oakland, California) (Coordinates CH201-14H23 for ZmME and CH201-117G14 for ZmMDH) by PCR using primers (5’-ACGACGGCCAGTGCCAAGCTTCCCTTCCGTCAGCAGATTAGGCG and 5’-ATTATTATGGAGAAACTCGAGGCAACATGGTTCTGGACCGATTCAG for ZmMDH; 5’-ACGACGGCCAGTGCCAAGCTTGGAATGACCACGAAATCGTCAAGCTAATCC and 5’-ATTATTATGGAGAAACTCGAGCTGTTACTGCTCTTTCCACTACTGAAGCAG for ZmME and sub-cloned into pSC0 vector. A binary vector with the hygromycin B resistance gene, pCAMBIA1300, was co-transformed with these vectors to allow for selection. To drive enriched ZmCA2 (GRMZM2G348512) expression in rice mesophyll cells, the vector of pSC110/ZmCA2-ACV5 was generated as previously described [60]. The rice transformation was performed at International Rice Research Institute (IRRI; Los Baños, Philippines) following a previously described method [57].

For ZmPEPC, ZmPPDK, ZmMDH and ZmME, after transformation, T_0_ PCR positive plants that had similar protein accumulation relative to the maize control were advanced to further generations to obtain homozygous lines with a single copy of the transgene. The selection of the four ZmPEPC events was described by [58]. These plants were used in the present study at the T_3_ generation for event ZmPEPC-60 and T4 generation for events ZmPEPC-28, ZmPEPC-62 and ZmPEPC-76. A total of 82 T_0_ plants were ZmPPDK PCR positive and seven plants (events) among them had a single copy of the transgene insertion. Four events with single transgene insertion and similar levels of ZmPPDK protein accumulation relative to the maize control were advanced to further generation to obtain homozygous lines. At T_2_ generation, three events (ZmPPDK-2, ZmPPDK-11 and ZmPPDK-52) were homozygous and used for analysis in the present study. A total of 37 T_0_ plants were ZmMDH PCR positive and eight plants among them had more than 50% ZmMDH protein accumulation compared to the maize control. However, all of them had multiple copies of the transgene insertion. From the segregating population at the T_1_ generation, we obtained plants with a single copy of the transgene from four events. Three events with a single copy of the transgene (ZmMDH-22, ZmMDH-43, and ZmMDH-48) were homozygous at T_2_ generation. The plants used in this study were at T_2_ generation for event ZmMDH-22 and T_3_ generation for events ZmMDH-43 and ZmMDH-48. A total of 46 T_0_ plants were PCR positive for the ZmME transgene and only two of them showed detectable ZmME protein accumulation by western blot analysis. Both events had more than six copies of the transgene. We obtained one homozygous event with seven copies of the transgene (ZmME-116) at T_2_ generation and used its T_3_ progeny in the present study. For ZmCA2, a total of 112 T_0_ plants were ZmCA2 PCR positive and eight plants (events) with a single copy of the transgene and highest abundant ZmCA2 accumulation were advanced to further generations to obtain homozygous lines. Three ZmCA2 events (ZmCA2-18, ZmCA2-39 and ZmCA2-69) were homozygous at T_2_ generation and used in the present study. The homozygosity of the transgene was confirmed by DNA blot analysis for T-DNA insertion (Supplemental Figure S1). The abundance of C_4_ protein accumulation was detected by western blot analysis (Supplemental Figure S2).

### Leaf chlorophyll content and plant growth analysis

Measurements of leaf chlorophyll content, tiller number and plant height were taken from plants at maximum tillering stage. The plants were grown in rice paddy fields at IRRI. Leaf chlorophyll content was measured in the upper youngest fully expanded leaves using a SPAD chlorophyll Meter (SPAD, Konica Minolta). Chlorophyll SPAD values are the average ± SE of three leaves from 15 plants per line. Tiller number and plant height are the average ± SE from 15 plants per line.

### Immunofluorescence microscopy

The middle portion of the seventh fully expanded leaf was sampled between 09:00 h and 11:00 h from 9-week-old plants. Leaf sections for immunolocalization analysis were prepared as described previously [57]. For detecting ZmCA2-AcV5 protein, the fixed sections were probed with the anti-AcV5 tag primary mouse monoclonal antibody (Abcam, Cambridge, UK) and the Alexa Fluor 488 goat anti-mouse IgG (Invitrogen) secondary antibody. For detecting ZmPPDK protein, the fixed sections were probed with the anti-PPDK primary rabbit polyclonal antibody (provided by Dr. Chris Chastain, Minnesota State University-Moorhead) and the Alexa Fluor 488 goat anti-rabbit IgG (Invitrogen) secondary antibody. The sections were examined on a BX61 with Disk Scanning Unit attachment microscope (Olympus) with florescence functions.

### RNA isolation, sequencing, and differential expression analysis

Total RNA was extracted using Trizol extraction methods (Invitrogen) and treated with RQ1 RNAase free DNAase (Promega) following a previously described method [57]. Leaf samples were harvested between 10:30 h and 11:30 h from 2-week-old plants. Total RNA was extracted from a pool of 12 fifth youngest fully expanded leaves from 12 individual plants for each line. For ME-116 and PEPC-28 lines, we extracted 3 individual RNA pools and each pool still contained 12 fifth leaves from 12 individual plants. RNA quality and quantity was checked using a NanoDrop ND-8000 spectrophotometer (Thermo Scientific, Waltham, MA, USA) and agarose gel electrophoresis. RNA samples were sequenced using Illumina platforms at The Beijing Genome Institute (BGI Tech Solutions (Hongkong) CO, Limited, Shenzhen, China). The reads were paired end (PE) of size 100 base pairs and the total amount of data was 4 Giga-bases of genomic sequence per sample. RNA-Seq reads were deposited to EBI array express and are available under the accession number E-MTAB-8539.

Raw reads were subject to quality trimming using TRIMMOMATIC [61]. This was done to remove low quality bases and read-pairs as well as contaminating adaptor sequences prior to transcript quantitation. Sequences were searched for all common Illumina adaptors (the default option) and the settings used for read processing by trimmomatic were LEADING:10 TRAILING:10 SLIDINGWINDOW:5:15 MINLEN:50. Quality trimmed reads were mapped to the full set of predicted transcript sequences from the *Oryza sativa* genome reference (323 v7) obtained from Phytozome version 12 [62] using Salmon [63]. Correlation in genome-wide transcript abundance estimates are shown in Supplemental Figure S5. Transcript abundance counts were summed at the locus level and differentially expressed transcripts were identified using DESeq2 as those genes with an adjusted *p*-value of < 0.01 [64].

### mRNA variance estimation in *Arabidopsis thaliana* natural accessions

The transcript abundance estimates for 17 different *Arabidopsis thaliana* natural accessions was downloaded from EBI array express under the accession number E-GEOD-53197. The floral bud and root samples were discarded and the standard deviation of the mRNA abundance estimates for each gene was calculated from all 17 aerial part samples. The percentile rank of each gene was then calculated. The full dataset is provided in Supplemental File S1.

### Phylogenetic analysis

The predicted protein sequences corresponding to the primary transcripts of 42 sequenced plant genomes were obtained from Phytozome v10 [62]. OrthoFinder [65,66] was used to infer orthogroups from these protein sequences. MAFFT-LINSI [67] was used to create multiple sequence alignments of the proteins within each orthogroup, and FastTree [68] was used to infer maximum likelihood phylogenetic trees from these multiple sequence alignments. The *Arabidopsis thaliana* orthologs of the maize genes used in the transgenic lines were identified using these phylogenetic trees (Supplemental File S3). If multiple paralogous genes existed in *Arabidopsis thaliana,* then the paralog that was recruited to C_4_ function in the closest C_4_ relative of *Arabidopsis thaliana*, *Gynandropsis gynandra* [10,69,70] was used for the analysis presented in Figure 2. The complete expression dataset for all paralogs is provided in Supplemental File S1.

### Immunoblot analyses

The middle portion of the seventh fully expanded leaf was sampled between 09:00 h and 11:00 h from 9-week-old plants. Soluble proteins were extracted and fractionated as described previously [57,58]. Samples were loaded based on equal leaf area (0.394 mm^2^ for ZmPEPC and ZmPPDK, and 3.94 mm^2^ for ZmMDH, ZmME and ZmCA2-AcV5). Proteins were electroblotted onto a polyvinylidene difluoride membrane and probed with antisera against AcV5 tag (Abcam, Cambridge, UK), ZmPPDK (provide by Chris Chastain, Minnesota State University, USA), ZmPEPC, ZmMDH or ZmME (provided by Richard Leegood, Sheffield University, UK) protein. The dilutions of ZmPEPC, ZmPPDK, ZmMDH, ZmME and AcV5 antibodies were 1:20000, 1:20000, 1:5000, 1:5000 and 1:2000, respectively. A peroxidase-conjugated secondary antibody was used at a dilution of 1:5000 and immunoreactive bands were visualized with ECL Western Blotting Detection Reagents (GE Healthcare, UK).

### Transgene insertion copy number estimation

Genomic DNA was extracted from leaves of matured plants and prepared for DNA blot analysis as described previously [57]. Genomic DNA of ZmCA2 lines at T_2_ generation was digested with BglII restriction endonuclease (NEB) and probed with a ZmPEPC promoter-specific probe as described previously [57,58]. Genomic DNA of ZmMDH and ZmPPDK lines at T_2_ generation was digested with EcoRI (NEB) and probed with their gene-specific probes generated from primers (5’-GAACCGCCAGAGTAGCAGAC and 5’-ATCGACGTATACGGCTGGTC for ZmMDH; 5’-TGTGGCGCCATGTTAGATAG and 5’-AATTCGTGAACACCCAGACC for ZmPPDK). The copy of number of ZmME insertion was detected by digesting genomic DNA with BamH1 (NEB) and probing with ZmME specific probe synthesized using primers (5’-TGGAGCTGCTTCCTTTTGTT and 5’-TGATAGGCAAGCACTGCAAC).

### Enzyme assays

Leaf samples for enzyme activity assay were harvested between 0900 and 1100 h from the youngest fully-expanded leaf of 4 to 5 –week-old plants. For ZmPEPC, ZmPPDK, ZmMDH and ZmME lines, proteins were extracted by homogenizing leaf material in a 250 μl of extraction buffer containing 50 mM Hepes-KOH, pH7.4, 5 mM MgCl_2_, 1 mM EDTA, 1 mM DTT, 1% (v/v) glycerol. After centrifugation at 10,000g for 2 minutes at 4°C, the supernatant was collected for enzyme activity assays. The PEPC enzyme activity assay was performed following previous methods [71,72]. The PEPC reaction mixture contained 100 mM Hepes-NaOH, pH 7.5, 10 mM MgCl_2_, 1 mM NaHCO3, 5 mM G6P, 0.2 mM NADH, 12 unit/ml MDH (from pig heart; Roche Diagnostics, Basel) and 4 mM PEP, and the reaction was started by adding PEP. The PPDK enzyme activity assay was performed as described by [36]. The MDH activity was determined by the method modified from [73]. The MDH reaction mixture contained 50 mM Hepes-KOH, pH 8, 70 mM KCl, 1 mM EDTA, 1 mM DTT, 1 mM OAA and 0.2 mM NADPH, and the reaction was started by adding OAA. The ME activity was measured by the method modified from [73]. The activities of PEPC, PPDK, MDH and ME were measured spectrophotometrically at 340 nm at 25°C, 30°C, 25°C and 25°C, respectively. The CA enzyme activity assay was conducted following a published method [74] using the electrometric method [75]. Proteins were extracted by homogenizing the leaf materials in a 250 μl of extraction buffer containing 50 mM Hepes-KOH, pH 7.2, 1 mM EDTA and 10 mM DTT. After centrifugation, the supernatant was collected for CA enzyme activity assay. The CA reaction buffer contained 15 mM Sodium Barbital and 3 mM Barbital; pH 8.3. All buffers and the assay were stored and performed on ice. The reaction was initiated by adding 8 ml of ice-cold CO_2_ saturated water into a reaction mixture containing 200 μl of protein extract and 12 ml of reaction buffer. The time required for a pH drop from 8.3 to 6.3 was recorded. The CA activity units were calculated as described previously [74].

### Photosynthesis assays

Leaf gas-exchange measurements were made at IRRI (mean atmospheric pressure of 94.8 kPa) using a LI-6400XT portable photosynthesis system (LICOR Biosciences, Lincoln, NE, USA). The system set–up was as previously described [57]. In summary, mmeasurements were taken at a constant airflow rate of 400 μmol s^−1^, leaf temperature of 30 °C and a leaf-to-air vapour pressure deficit of between 1.5 and 2 kPa. Data was acquired between 0800 h and 1300 h in a room with an air temperature maintained at approximately 30 °C. Measurements were made on the mid-portion of the leaf blade of three fully expanded leaves during the tillering stage for each transgenic event from two to three plants from each transgenic line. Leaves were acclimated in the cuvette for approximately 30 min before measurements were made. The response curves of the net rate of CO_2_ assimilation (A, μmol CO_2_ m^−2^ s^−1^) to changing intercellular CO_2_ concentration (C*i*, μmol CO_2_ mol^−1^) were acquired by increasing *C_a_* (CO_2_ concentration in the cuvette) from 20 to 2,000 μmol CO_2_ mol^−1^ at a photosynthetic photon flux density (PPFD) of 2,000 μmol photons m^−2^ s^−1^.

## Acknowledgements

This work was funded by the Bill & Melinda Gates Foundation through award number OPP1129902. SK is a Royal Society University Research Fellow. Work in SKs lab is supported by the Royal Society and the European Union’s Horizon 2020 research and innovation program under grant agreement number 637765. Technical assistance was provided by Justina Davis, Jean Melgar and Flor Montecillo for rice transfomation, tissue culture and handling of the transgnic plants.

## Supplemental Figures

**Supplemental File S1**

Excel spreadsheet. This spreadsheet contains the transcripts per million (TPM) estimates of relative mRNA abundance for the aerial part of all 17 natural accessions of *Arabidopsis thaliana*. The standard deviation and percentile are also provided. For convenience the genes of relevance are provided at the top of the spreadsheet.

**Supplemental File S2**

Online isotope discrimination analysis of two transgenic lines overexpressing ZmCA.

**Supplemental File S3**

Excel Spreadsheet. This spreadsheet contains the transcripts per million (TPM) estimates of relative mRNA abundance for all genes in the transgenic rice lines overexpressing core C_4_ cycle enzymes. It also contains the differentially expressed gene lists.

**Supplemental File S4**

PDF. This file contains the phylogenetic trees used to identify *Arabidopsis thaliana* orthologs of the maize genes used in this study.

---

**Supplemental Figure S1.**
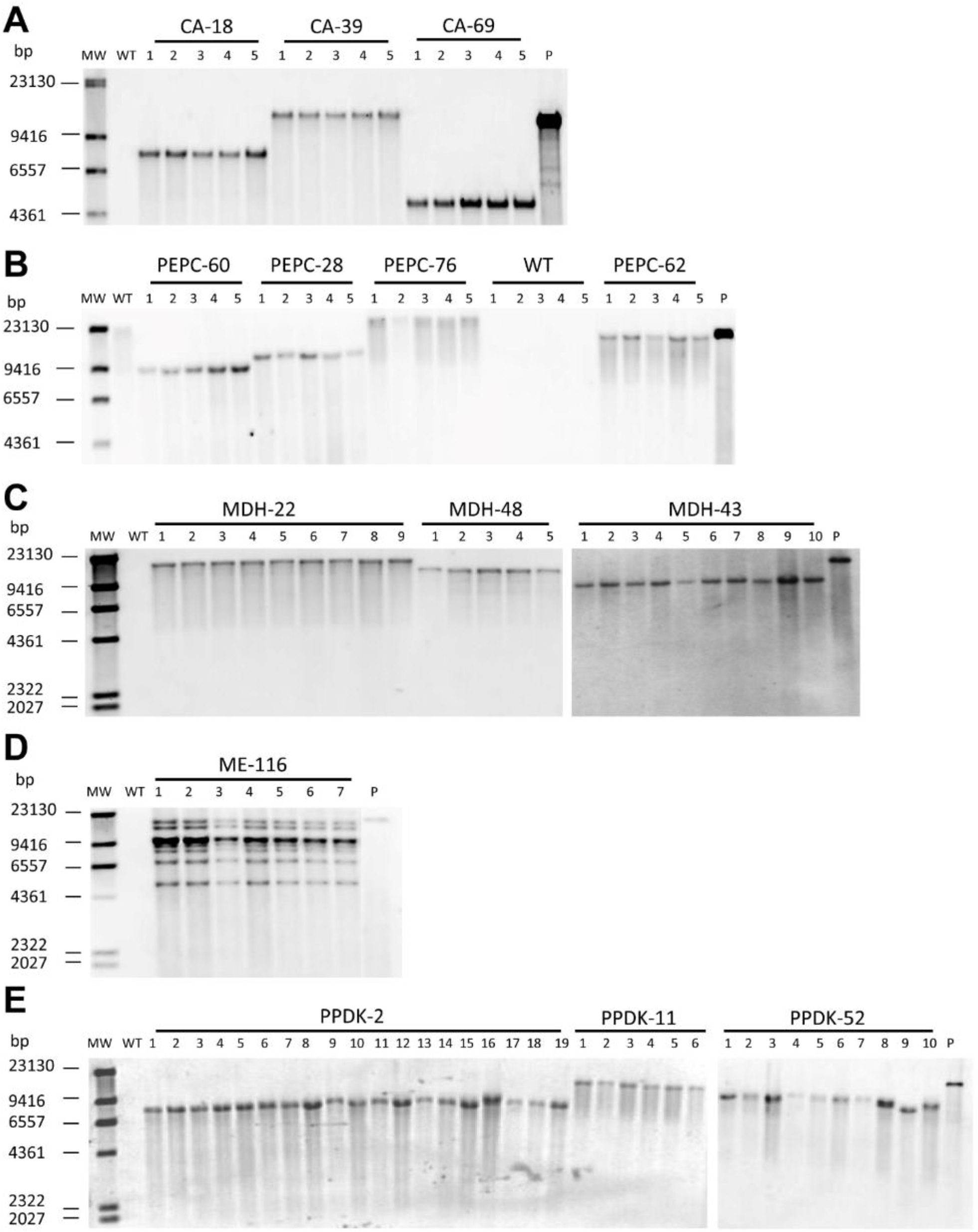
Southern blot analysis of insertion copy number in the transgenic lines. In all cases DNA isolated from different individual plants descendant from the same transgenic event are shown. The name and number of the independent transgenic event is shown on the top line and the individual plant number is shown below. A) Three independent single insertion lines containing the construct for overexpression of *Zea mays* β-carbonic anhydrase 2 (CA-18, CA-39 and CA-69). B) Four independent single insertion lines containing the construct for overexpression of *Zea mays* PEPC (PEPC-28, PEPC-60, PEPC-62 and PEPC-76). C) Three independent single insertion lines for overexpression of *Zea mays* malate dehydrogenase (MDH-22, MDH-43 and MDH-48). D) A single transgenic event containing multiple insertions (≥6) of the construct for overexpression of *Zea mays* malic enzyme (ME-116). E) Three independent single insertion lines for overexpression of the *Zea mays* PPDK (PPDK-2, PPDK-11 and PPDK-52). In all cases WT corresponds to genomic DNA isolated from untransformed wild type rice plants. The un-digested plasmid is used as a positive control (P). Sizes of molecular weight markers are indicated next to the image.

**Supplemental Figure S2.**
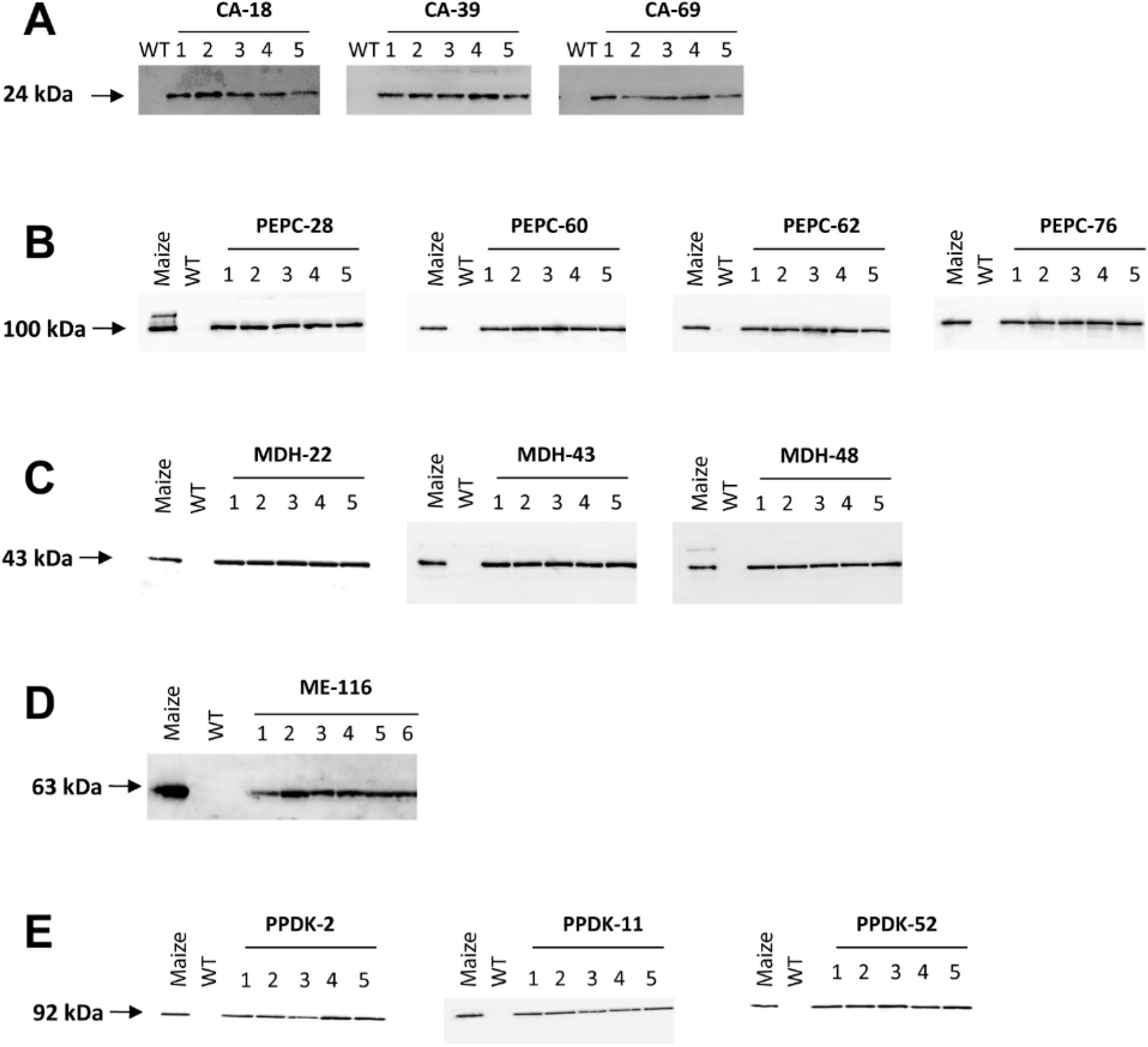
Western blot analysis of protein expression in transgenic lines. A) Detection of *Zea mays* β-carbonic anhydrase 2 protein expression in multiple plants descendent from the three independent transgenic events. Protein expression was detected using the anti-AcV5 antibody as the CDS of the ZmCA gene was modified to contain an Ac-V5 epitope tag at the C-terminus of the protein. B) Detection of *Zea mays* phosphoenolpyruvate carboxylase (ZmPEPC) protein expression in multiple plants descendent from the four independent transgenic events. Protein expression was detected using the PEPC antibody. Protein isolated from wild type maize plants (maize) was used as a positive control, while protein isolated from wild-type rice plants (WT) was used as a negative control. C) Detection of *Zea mays* malate dehydrogenase (ZmMDH) protein expression in multiple plants descendent from the three independent transgenic events. Protein expression was detected using the MDH antibody. Protein isolated from wild type maize plants (maize) was used as a positive control, while protein isolated from wild-type rice plants (WT) was used as a negative control. D) Detection of *Zea mays* malic enzyme (ZmME) protein expression in multiple plants descendent from the single transgenic event used in this study. Protein expression was detected using the ME antibody. Protein isolated from wild type maize plants (maize) was used as a positive control, while protein isolated from wild-type rice plants (WT) was used as a negative control. E) Detection of *Zea mays* pyruvate, phosphate dikinase (ZmPPDK) protein expression in multiple plants descendent from the three independent transgenic events. Protein expression was detected using the PPDK antibody. Protein isolated from wild type maize plants (maize) was used as a positive control, while protein isolated from wild-type rice plants (WT) was used as a negative control.

**Figure S3.**
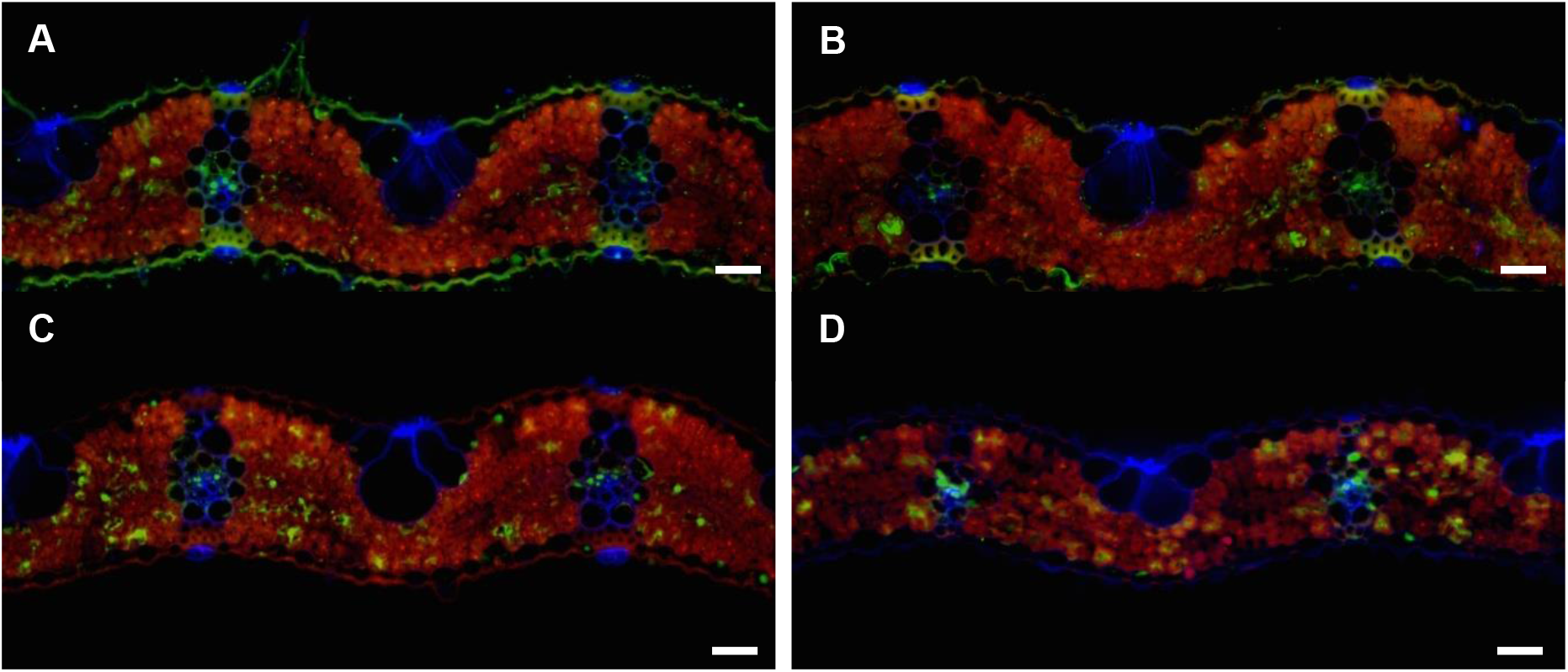
Localisation of overexpressed proteins. A & B) Rice leaf cross sections stained with anti-MDH antibody for A) an un-transformed wild type plant and B) a transgenic line overexpressing ZmMDH. C & D) Rice leaf cross sections stained with anti-ZmME antibody for C) an un-transformed wild-type plant and D) a transgenic line overexpression ZmME. Magnification: 200x. Scale bar: 20 μm.

**Supplemental Figure S4.**
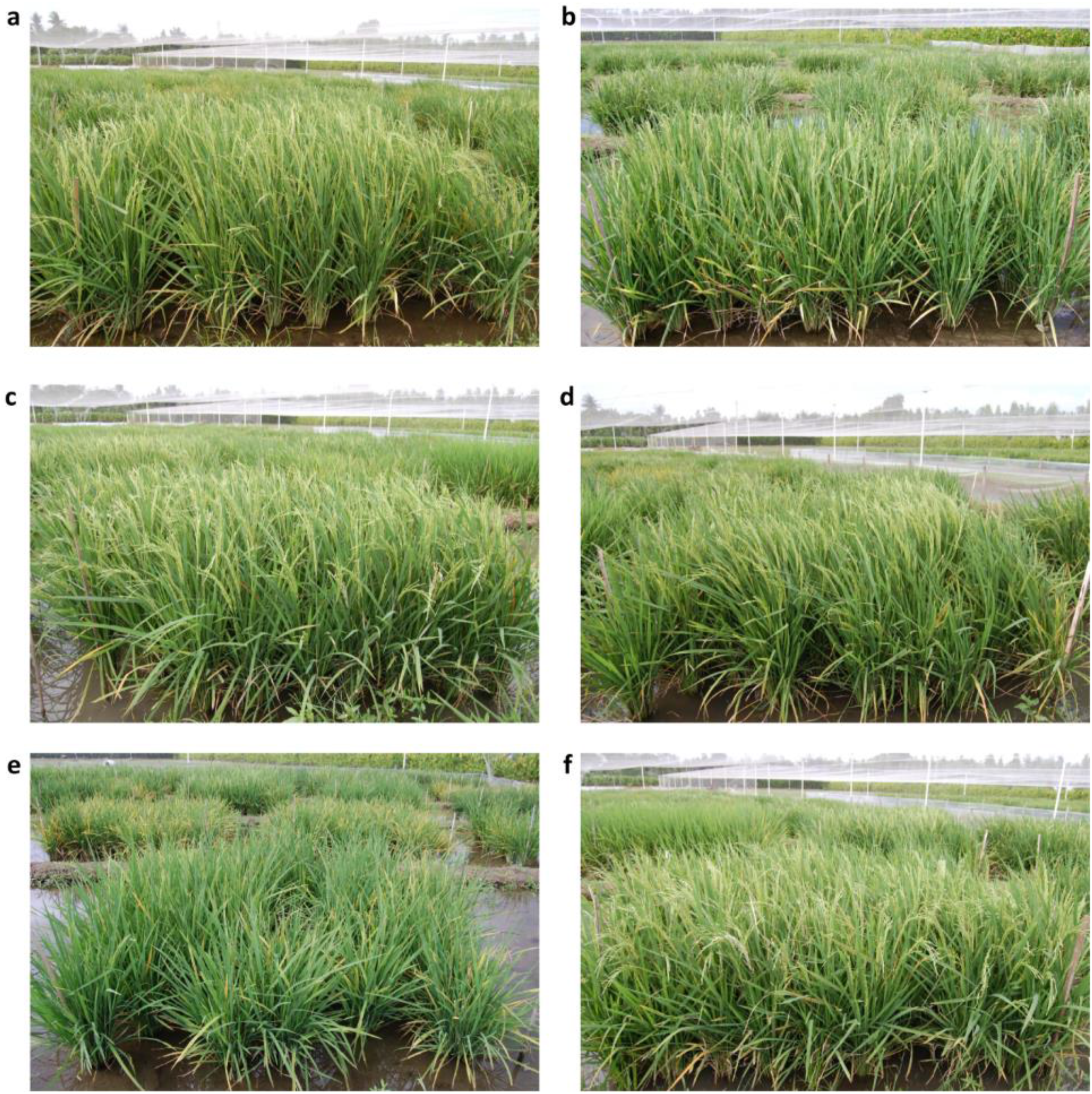
Representative images of the transgenic lines growing in field conditions at 80 days post germination. A) Wild type rice plants. B) Transgenic line CA-39. C) Transgenic line PEPC-76. D) Transgenic line MDH-48. E) Transgenic line ME-116. F) Transgenic line PPDK-52.

**Supplemental Figure S5.**
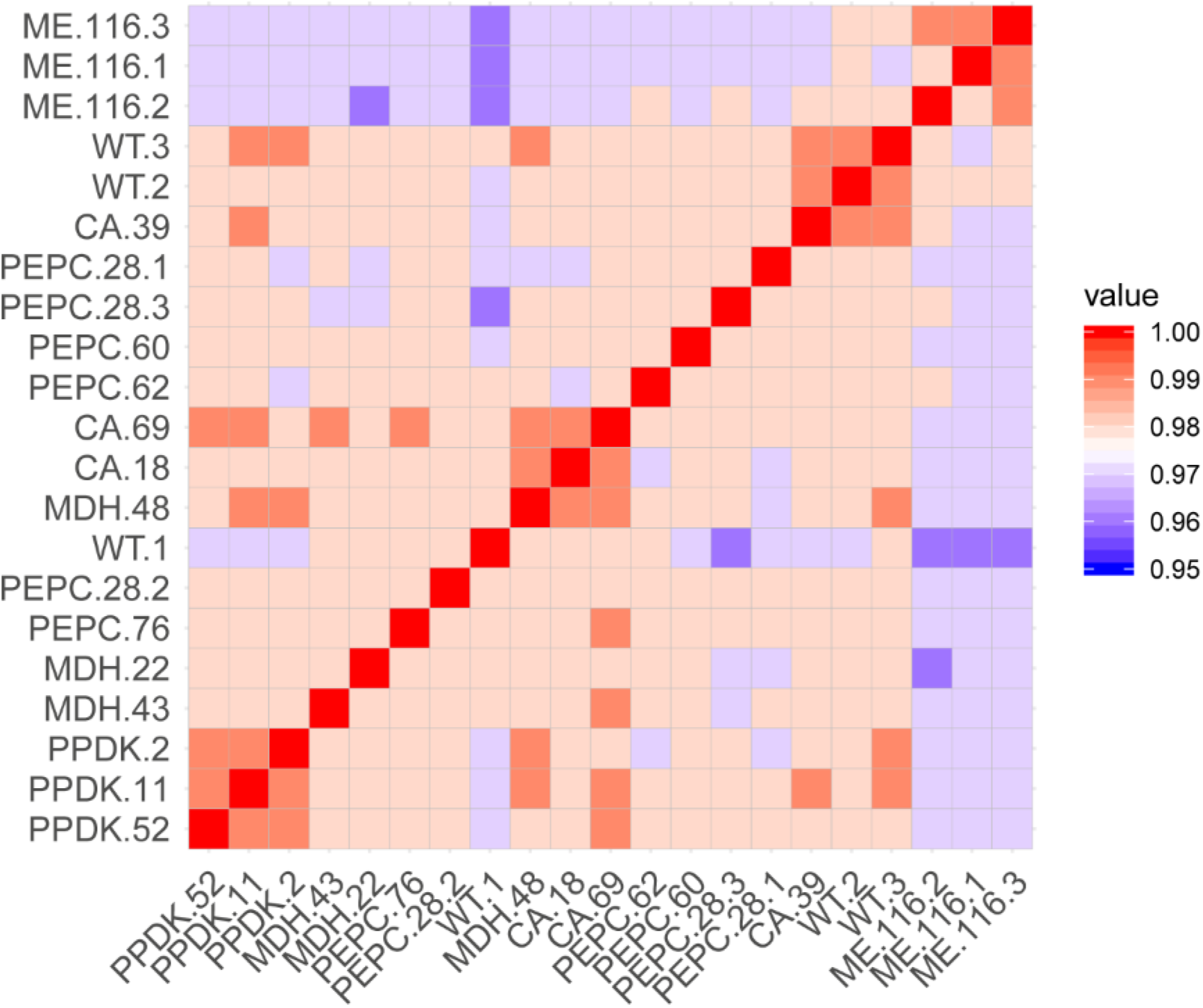
Correlation in genome-wide transcript abundance estimates between samples. Only genes that were detected as expressed in all samples were used in to inform the correlation plot. Relative mRNA abundance estimates (transcripts per million) were log transformed and used to calculate pairwise Pearson correlation coefficients. Correlation plot shows that mRNA abundance estimates in transgenic lines expressing ZmME are more similar to each other then they are to other samples, consistent with the larger number of differentially expressed genes identified in these lines. All other samples are broadly very similar, consistent with the lack of growth or photosynthesis phenotypes observed in these plants.

## References

1. Stern DL: The genetic causes of convergent evolution. Nature Reviews Genetics 2013, 14:751–764.

2. Losos JB: CONVERGENCE, ADAPTATION, AND CONSTRAINT. Evolution 2011, 65:1827–1840.

3. Tokita M: How the pterosaur got its wings. Biological Reviews 2015, 90:1163–1178.

4. Schwab IR: The evolution of eyes: major steps. The Keeler lecture 2017: centenary of Keeler Ltd. Eye 2018, 32:302–313.

5. Kozmik Z, Ruzickova J, Jonasova K, Matsumoto Y, Vopalensky P, Kozmikova I, Strnad H, Kawamura S, Piatigorsky J, Paces V, et al.: Assembly of the cnidarian camera-type eye from vertebrate-like components. Proc Natl Acad Sci U S A 2008, 105:8989–8993.

6. Parker J, Tsagkogeorga G, Cotton JA, Liu Y, Provero P, Stupka E, Rossiter SJ: Genome-wide signatures of convergent evolution in echolocating mammals. Nature 2013, 502:228–231.

7. Manceau M, Domingues VS, Linnen CR, Rosenblum EB, Hoekstra HE: Convergence in pigmentation at multiple levels: mutations, genes and function. Philosophical transactions of the Royal Society of London. Series B, Biological sciences 2010, 365:2439–2450.

8. Bräutigam A, Kajala K, Wullenweber J, Sommer M, Gagneul D, Weber KL, Carr KM, Gowik U, Maß J, Lercher MJ, et al.: An mRNA Blueprint for C<sub>4</sub> Photosynthesis Derived from Comparative Transcriptomics of Closely Related C<sub>3</sub> and C<sub>4</sub> Species. Plant Physiology 2011, 155:142–156.

9. Gowik U, Bräutigam A, Weber KL, Weber APM, Westhoff P: Evolution of C<sub>4</sub> Photosynthesis in the Genus <em>Flaveria</em>: How Many and Which Genes Does It Take to Make C<sub>4</sub>? The Plant Cell 2011, 23:2087–2105.

10. Kelly S, Covshoff S, Wanchana S, Thakur V, Quick WP, Wang Y, Ludwig M, Bruskiewich R, Fernie AR, Sage RF, et al.: Wide sampling of natural diversity identifies novel molecular signatures of C<sub>4</sub> photosynthesis. bioRxiv 2017:163097.

11. Külahoglu C, Denton AK, Sommer M, Maß J, Schliesky S, Wrobel TJ, Berckmans B, Gongora-Castillo E, Buell CR, Simon R, et al.: Comparative Transcriptome Atlases Reveal Altered Gene Expression Modules between Two Cleomaceae C<sub>3</sub> and C<sub>4</sub> Plant Species. The Plant Cell 2014, 26:3243–3260.

12. Sedelnikova OV, Hughes TE, Langdale JA: Understanding the Genetic Basis of C4 Kranz Anatomy with a View to Engineering C3 Crops. Annual Review of Genetics 2018, 52:249–270.

13. Sage RF: A portrait of the C4 photosynthetic family on the 50th anniversary of its discovery: species number, evolutionary lineages, and Hall of Fame. J Exp Bot 2016, 67:4039–4056.

14. Zhu X-G, Long SP, Ort DR: What is the maximum efficiency with which photosynthesis can convert solar energy into biomass? Current Opinion in Biotechnology 2008, 19:153–159.

15. Brown RH: A Difference in N Use Efficiency in C3 and C4 Plants and its Implications in Adaptation and Evolution1. Crop Science 1978, 18:93–98.

16. Sage RF, Pearcy RW, Seemann JR: The Nitrogen Use Efficiency of C<sub>3</sub> and C<sub>4</sub> Plants. <span class=“subtitle”>III. Leaf Nitrogen Effects on the Activity of Carboxylating Enzymes in <em>Chenopodium album</em> (L.) and <em>Amaranthus retroflexus</em> (L.)</span> 1987, 85:355–359.

17. von Caemmerer S, Furbank RT: Strategies for improving C4 photosynthesis. Current Opinion in Plant Biology 2016, 31:125–134.

18. Edwards EJ, Osborne CP, Stromberg CA, Smith SA, Consortium CG, Bond WJ, Christin PA, Cousins AB, Duvall MR, Fox DL, et al.: The origins of C4 grasslands: integrating evolutionary and ecosystem science. Science 2010, 328:587–591.

19. Hibberd JM, Sheehy JE, Langdale JA: Using C4 photosynthesis to increase the yield of rice-rationale and feasibility. Curr Opin Plant Biol 2008, 11:228–231.

20. von Caemmerer S, Quick WP, Furbank RT: The development of C(4)rice: current progress and future challenges. Science 2012, 336:1671–1672.

21. Huang P, Brutnell TP: A synthesis of transcriptomic surveys to dissect the genetic basis of C4 photosynthesis. Curr Opin Plant Biol 2016, 31:91–99.

22. Niklaus M, Kelly S: The molecular evolution of C4 photosynthesis: opportunities for understanding and improving the world's most productive plants. J Exp Bot 2019, 70:795–804.

23. Osborne CP, Sack L: Evolution of C4 plants: a new hypothesis for an interaction of CO2 and water relations mediated by plant hydraulics. Philos Trans R Soc Lond B Biol Sci 2012, 367:583–600.

24. Christin P-A, Osborne CP: The evolutionary ecology of C4 plants. New Phytologist 2014, 204:765–781.

25. Brown NJ, Aubry S, Hibberd JM: The role of proteins in C3 plants prior to their recruitment into the C4 pathway. Journal of Experimental Botany 2011, 62:3049–3059.

26. Ludwig M: Evolution of carbonic anhydrase in C4 plants. Curr Opin Plant Biol 2016, 31:16–22.

27. Gowik U, Westhoff P: The Path from C<sub>3</sub> to C<sub>4</sub> Photosynthesis. Plant Physiology 2011, 155:56–63.

28. Sedelnikova OV, Hughes TE, Langdale JA: Understanding the Genetic Basis of C4 Kranz Anatomy with a View to Engineering C3 Crops. Annu Rev Genet 2018, 52:249–270.

29. Christin PA, Besnard G, Samaritani E, Duvall MR, Hodkinson TR, Savolainen V, Salamin N: Oligocene CO2 decline promoted C4 photosynthesis in grasses. Curr Biol 2008, 18:37–43.

30. Feodorova TA, Voznesenskaya EV, Edwards GE, Roalson EH: Biogeographic Patterns of Diversification and the Origins of C_4_ in Cleome (Cleomaceae). Systematic Botany 2010, 35:811–826.

31. Heckmann D, Schulze S, Denton A, Gowik U, Westhoff P, Weber AP, Lercher MJ: Predicting C4 photosynthesis evolution: modular, individually adaptive steps on a Mount Fuji fitness landscape. Cell 2013, 153:1579–1588.

32. Williams BP, Johnston IG, Covshoff S, Hibberd JM: Phenotypic landscape inference reveals multiple evolutionary paths to C4 photosynthesis. Elife 2013, 2:e00961.

33. Sage RF: The evolution of C4 photosynthesis. New Phytologist 2004, 161:341–370.

34. Blätke M-A, Bräutigam A: Evolution of C4 photosynthesis predicted by constraint-based modelling. eLife 2019, 8:e49305.

35. Gibson G, Dworkin I: Uncovering cryptic genetic variation. Nat Rev Genet 2004, 5:681–690.

36. Le Rouzic A, Carlborg O: Evolutionary potential of hidden genetic variation. Trends Ecol Evol 2008, 23:33–37.

37. Paaby AB, Rockman MV: Cryptic genetic variation: evolution's hidden substrate. Nat Rev Genet 2014, 15:247–258.

38. Amitai G, Gupta RD, Tawfik DS: Latent evolutionary potentials under the neutral mutational drift of an enzyme. HFSP J 2007, 1:67–78.

39. Bloom JD, Romero PA, Lu Z, Arnold FH: Neutral genetic drift can alter promiscuous protein functions, potentially aiding functional evolution. Biol Direct 2007, 2:17.

40. de Visser JA, Cooper TF, Elena SF: The causes of epistasis. Proc Biol Sci 2011, 278:3617–3624.

41. Queitsch C, Sangster TA, Lindquist S: Hsp90 as a capacitor of phenotypic variation. Nature 2002, 417:618–624.

42. Wagner A: Neutralism and selectionism: a network-based reconciliation. Nat Rev Genet 2008, 9:965–974.

43. Gan X, Stegle O, Behr J, Steffen JG, Drewe P, Hildebrand KL, Lyngsoe R, Schultheiss SJ, Osborne EJ, Sreedharan VT, et al.: Multiple reference genomes and transcriptomes for Arabidopsis thaliana. Nature 2011, 477:419–423.

44. Weigel D, Mott R: The 1001 Genomes Project for Arabidopsis thaliana. Genome Biology 2009, 10:107.

45. The rgp: The 3,000 rice genomes project. GigaScience 2014, 3:7.

46. Mackill DJ, Khush GS: IR64: a high-quality and high-yielding mega variety. Rice (N Y) 2018, 11:18.

47. Schomburg I, Jeske L, Ulbrich M, Placzek S, Chang A, Schomburg D: The BRENDA enzyme information system–From a database to an expert system. Journal of Biotechnology 2017, 261:194–206.

48. Denton AK, Mass J, Kulahoglu C, Lercher MJ, Brautigam A, Weber AP: Freeze-quenched maize mesophyll and bundle sheath separation uncovers bias in previous tissue-specific RNA-Seq data. J Exp Bot 2017, 68:147–160.

49. Chang YM, Liu WY, Shih AC, Shen MN, Lu CH, Lu MY, Yang HW, Wang TY, Chen SC, Chen SM, et al.: Characterizing regulatory and functional differentiation between maize mesophyll and bundle sheath cells by transcriptomic analysis. Plant Physiol 2012, 160:165–177.

50. Majeran W, Cai Y, Sun Q, van Wijk KJ: Functional differentiation of bundle sheath and mesophyll maize chloroplasts determined by comparative proteomics. Plant Cell 2005, 17:3111–3140.

51. Sasaki H, Hirose T, Watanabe Y, Ohsugi R: Carbonic anhydrase activity and CO2-transfer resistance in Zn-deficient rice leaves. Plant Physiol 1998, 118:929–934.

52. Taniguchi Y, Ohkawa H, Masumoto C, Fukuda T, Tamai T, Lee K, Sudoh S, Tsuchida H, Sasaki H, Fukayama H, et al.: Overproduction of C4 photosynthetic enzymes in transgenic rice plants: an approach to introduce the C4-like photosynthetic pathway into rice. J Exp Bot 2008, 59:1799–1809.

53. Takeuchi Y, Akagi H, Kamasawa N, Osumi M, Honda H: Aberrant chloroplasts in transgenic rice plants expressing a high level of maize NADP-dependent malic enzyme. Planta 2000, 211:265–274.

54. Evans JR, Von Caemmerer S: Temperature response of carbon isotope discrimination and mesophyll conductance in tobacco. Plant, Cell & Environment 2013, 36:745–756.

55. Gillon JS, Yakir D: Internal Conductance to CO<sub>2</sub> Diffusion and C<sup>18</sup>OO Discrimination in C<sub>3</sub> Leaves. Plant Physiology 2000, 123:201–214.

56. Taylor L, Nunes-Nesi A, Parsley K, Leiss A, Leach G, Coates S, Wingler A, Fernie AR, Hibberd JM: Cytosolic pyruvate,orthophosphate dikinase functions in nitrogen remobilization during leaf senescence and limits individual seed growth and nitrogen content. The Plant Journal 2010, 62:641–652.

57. Lin H, Karki S, Coe RA, Bagha S, Khoshravesh R, Balahadia CP, Ver Sagun J, Tapia R, Israel WK, Montecillo F, et al.: Targeted Knockdown of GDCH in Rice Leads to a Photorespiratory-Deficient Phenotype Useful as a Building Block for C4 Rice. Plant Cell Physiol 2016, 57:919–932.

58. Giuliani R, Karki S, Covshoff S, Lin HC, Coe RA, Koteyeva NK, Evans MA, Quick WP, von Caemmerer S, Furbank RT, et al.: Transgenic maize phosphoenolpyruvate carboxylase alters leaf-atmosphere CO2 and (13)CO2 exchanges in Oryza sativa. Photosynth Res 2019.

59. Matsuoka M: Structure, genetic mapping, and expression of the gene for pyruvate, orthophosphate dikinase from maize. J Biol Chem 1990, 265:16772–16777.

60. Osborn HL, Alonso-Cantabrana H, Sharwood RE, Covshoff S, Evans JR, Furbank RT, von Caemmerer S: Effects of reduced carbonic anhydrase activity on CO2 assimilation rates in Setaria viridis: a transgenic analysis. J Exp Bot 2017, 68:299–310.

61. Bolger AM, Lohse M, Usadel B: Trimmomatic: a flexible trimmer for Illumina sequence data. Bioinformatics 2014, 30:2114–2120.

62. Goodstein DM, Shu S, Howson R, Neupane R, Hayes RD, Fazo J, Mitros T, Dirks W, Hellsten U, Putnam N, et al.: Phytozome: a comparative platform for green plant genomics. Nucleic Acids Res 2012, 40:D1178–1186.

63. Patro R, Duggal G, Love MI, Irizarry RA, Kingsford C: Salmon provides fast and bias-aware quantification of transcript expression. Nat Methods 2017, 14:417–419.

64. Love MI, Huber W, Anders S: Moderated estimation of fold change and dispersion for RNA-seq data with DESeq2. Genome Biol 2014, 15:550.

65. Emms DM, Kelly S: OrthoFinder: solving fundamental biases in whole genome comparisons dramatically improves orthogroup inference accuracy. Genome Biol 2015, 16:157.

66. Emms DM, Kelly S: OrthoFinder: phylogenetic orthology inference for comparative genomics. Genome Biology 2019, 20:238.

67. Katoh K, Standley DM: MAFFT Multiple Sequence Alignment Software Version 7: Improvements in Performance and Usability. Molecular Biology and Evolution 2013, 30:772–780.

68. Price MN, Dehal PS, Arkin AP: FastTree 2 – Approximately Maximum-Likelihood Trees for Large Alignments. PLOS ONE 2010, 5:e9490.

69. Aubry S, Aresheva O, Reyna-Llorens I, Smith-Unna RD, Hibberd JM, Genty B: A Specific Transcriptome Signature for Guard Cells from the C<sub>4</sub> Plant <em>Gynandropsis gynandra</em>. Plant Physiology 2016, 170:1345–1357.

70. Aubry S, Kelly S, Kümpers BMC, Smith-Unna RD, Hibberd JM: Deep Evolutionary Comparison of Gene Expression Identifies Parallel Recruitment of Trans-Factors in Two Independent Origins of C4 Photosynthesis. PLOS Genetics 2014, 10:e1004365.

71. Meyer CR, Rustin P, Wedding RT: A simple and accurate spectrophotometric assay for phosphoenolpyruvate carboxylase activity. Plant Physiol 1988, 86:325–328.

72. Ueno Y, Hata S, Izui K: Regulatory phosphorylation of plant phosphoenolpyruvate carboxylase: role of a conserved basic residue upstream of the phosphorylation site. FEBS Lett 1997, 417:57–60.

73. Tsuchida H, Tamai T, Fukayama H, Agarie S, Nomura M, Onodera H, Ono K, Nishizawa Y, Lee BH, Hirose S, et al.: High level expression of C4-specific NADP-malic enzyme in leaves and impairment of photoautotrophic growth in a C3 plant, rice. Plant Cell Physiol 2001, 42:138–145.

74. Yu S, Zhang X, Guan Q, Takano T, Liu S: Expression of a carbonic anhydrase gene is induced by environmental stresses in rice (Oryza sativa L.). Biotechnol Lett 2007, 29:89–94.

75. Wilbur KM, Anderson NG: Electrometric and colorimetric determination of carbonic anhydrase. J Biol Chem 1948, 176:147–154.

