## Supplemental File S2 for "A role for neutral variation in the evolution of C_4_ photosynthesis"

**Supplemental File S2. Table A.** Leaf functional traits estimated on rice CA overexpressing lines and WT based on simultaneous leaf-atmosphere $H_2O$, $CO_2$, and $^{13}C^{18}OO$ exchange measurements under 20 and 2 % [$O_2$] and different PPFD. LI-COR leaf cuvette was set with $C_a$ at 380 $\mu$mol mol$^{-1}$ and PPFD at 1500, 500 and 200 $\mu$mol photons m$^{-2}$ s$^{-1}$. Leaf temperature was 25 °C and leaf-atmosphere VPD was maintained in the range of 1.0-1.5 kPa. Values are mean ± SE (n= 4). $A$ = net $CO_2$ assimilation rate; $g_s$= stomatal conductance to $CO_2$; $C_i$= intercellular $CO_2$ molar fraction; $C_a$ = atmospheric $CO_2$ molar fraction. The chloroplastic $CO_2$ molar fraction ($C_c$) was estimated using mean mesophyll conductance to $CO_2$ ($g_m$) for each Tmt. $\Delta^{13}C$ = net discrimination against $^{13}CO_2$. Significance (P< 0.05) of the effects of Plant type, Tmt and Plant type * Tmt interaction was evaluated by SAS Proc Mixed according to a (split plot without replicates) between and within subjects statistical design: between subjects are Plant types; within subjects are the four Tmts, subjects are the leaves. The significance of $O_2$ level at PPFD of 1500 $\mu$mol photons m$^{-2}$ s$^{-1}$ was evaluated by polynomial contrast (PC); the significance of PPFD levels at 2 % [$O_2$] was evaluated in terms of linear and quadratic polynomial orthogonal contrasts (POC) for unequal PPFD increments.

| Plant type | Tmt | [$O_2$] | PPFD | $A$ | $g_s$ | $C_i$ | $C_i/C_a$ | $C_c$ | $g_m$ | $\Delta^{13}C$ |
|---|---|---|---|---|---|---|---|---|---|---|
| | | (mmol mol$^{-1}$) | ($\mu$mol photons m$^{-2}$ s$^{-1}$) | ($\mu$mol $CO_2$ m$^{-2}$ s$^{-1}$) | (mol $CO_2$ m$^{-2}$ s$^{-1}$) | ($\mu$mol mol$^{-1}$) | | ($\mu$mol mol$^{-1}$) | (mol $CO_2$ m$^{-2}$ s$^{-1}$) | (‰) |
| CA-18 | 1 | 200 | 1500 | 21.8 ± 0.7 | 0.46 ± 0.05 | 311 ± 5 | 0.80 ± 0.01 | 224 ± 13 | 0.26 ± 0.03 | 17.7 ± 1.3 |
| | 2 | 20 | 1500 | 28.6 ± 0.8 | 0.42 ± 0.06 | 282 ± 8 | 0.73 ± 0.02 | 177 ± 21 | 0.30 ±0.04 | 15.7 ± 0.7 |
| | 3 | 20 | 500 | 17.1 ± 1.0 | 0.17 ± 0.02 | 260 ± 10 | 0.67 ± 0.02 | 166 ± 18 | 0.20 ± 0.03 | 20.0 ± 1.3 |
| | 4 | 20 | 200 | 9.6 ± 0.5 | 0.12 ± 0.02 | 278 ± 10 | 0.71 ± 0.02 | 165 ± 15 | 0.09 ± 0.01 | 20.1 ± 1.4 |
| CA-39 | 1 | 200 | 1500 | 22.4 ± 0.3 | 0.44 ± 0.04 | 309 ± 4 | 0.79 ± 0.01 | 221 ± 12 | 0.27 ± 0.03 | 19.4 ± 1.2 |
| | 2 | 20 | 1500 | 28.8 ± 1.6 | 0.33 ± 0.05 | 267 ± 7 | 0.69 ± 0.02 | 167 ± 18 | 0.31 ± 0.04 | 14.9 ± 1.0 |
| | 3 | 20 | 500 | 19.7 ± 0.3 | 0.19 ± 0.01 | 259 ± 5 | 0.67 ± 0.02 | 179 ± 7 | 0.26 ± 0.03 | 18.9 ± 0.6 |
| | 4 | 20 | 200 | 11.8 ± 0.4 | 0.17 ± 0.01 | 294 ± 5 | 0.75 ± 0.02 | 215 ± 15 | 0.16 ± 0.02 | 21.1 ± 1.5 |
| WT | 1 | 200 | 1500 | 21.9 ± 1.1 | 0.46 ± 0.04 | 313 ± 4 | 0.80 ± 0.01 | 252 ± 6 | 0.34 ± 0.01 | 20.2 ± 1.6 |
| | 2 | 20 | 1500 | 24.7 ± 1.1 | 0.26 ± 0.01 | 261 ± 4 | 0.67 ± 0.01 | 153 ± 17 | 0.23 ± 0.02 | 13.1 ± 1.7 |
| | 3 | 20 | 500 | 17.4 ± 0.6 | 0.16 ± 0.01 | 253 ± 6 | 0.65 ± 0.02 | 177 ± 18 | 0.24 ± 0.04 | 16.8 ± 2.1 |
| | 4 | 20 | 200 | 10.3 ± 0.6 | 0.12 ± 0.01 | 277 ± 10 | 0.71 ± 0.02 | 193 ± 24 | 0.13 ± 0.02 | 18.3 ± 2.3 |
| | *Significance* | | *Plant type* | P= 0.03 | P= 0.54 | P= 0.73 | P= 0.75 | P= 0.62 | P= 0.57 | P= 0.32 |
| | | | *Tmt* | P< 0.0001 | P< 0.0001 | P< 0.0001 | P< 0.0001 | P< 0.0001 | P< 0.0001 | P= 0.0002 |
| | | | *Plant type * Tmt* | P= 0.19 | P= 0.16 | P= 0.28 | P= 0.29 | P= 0.24 | P= 0.40 | P= 0.50 |
| | | *Tmts 1-2* | *PC* | P< 0.0001 | P= 0.14 | P< 0.0001 | P< 0.0001 | P= 0.17 | P= 0.0001 | P= 0.28 |
| | | *Tmts 2-3-4* | *POC linear* | P< 0.0001 | P< 0.0001 | P< 0.0001 | P< 0.0001 | P= 0.001 | P= 0.32 | P= 0.42 |
| | | | *POC quadratic* | P< 0.0001 | P= 0.05 | P< 0.0001 | P< 0.0001 | P< 0.0001 | P= 0.90 | P= 0.0001 |

**Supplemental File S2. Table B.** Leaf functional traits estimated on rice CA overexpressing lines and WT based on simultaneous leaf-atmosphere $H_2O$, $CO_2$, $^{13}C^{18}OO$ and $H_2{}^{18}O$ exchange measurements under 20 and 2 % [$O_2$] and different PPFD. LI-COR leaf cuvette was set with $C_a$ at 380 $\mu$mol mol$^{-1}$ and PPFD at 1500, 500 and 200 $\mu$mol photons m$^{-2}$ s$^{-1}$. Leaf temperature was 25 °C and leaf-atmosphere VPD was maintained in the range of 1.0-1.5 kPa. Values are mean ± SE (n= 4). $\Delta^{18}O$= net discrimination against $C^{18}OO$; $\Delta_{ca}$= $C^{18}OO$ enrichment at the sites of oxygen exchange compared to atmosphere; $\Delta_{ea}$ = $C^{18}OO$ enrichment at the sites of oxygen exchange in full isotope equilibrium with water, compared to atmosphere; $\delta_t$ = $^{18}O$ composition of transpired water vapor; $\delta_e$ = $^{18}O$ composition of water at the sites of evaporation within the leaves; $\theta$ = Proportion of $CO_2$ in isotopic equilibrium with water at the sites of oxygen exchange. Correction for ternary effects, and correction for $O_2$ and humidity levels on $H_2{}^{18}O$ readings were performed. Significance ($P< 0.05$) of the effects of Plant type, Tmt and Plant type * Tmt interaction was evaluated by SAS Proc Mixed according to a (split plot without replicates) between and within subjects statistical design: between subjects are Plant types; within subjects are the four Tmts, subjects are the leaves. The significance of $O_2$ level at PPFD of 1500 $\mu$mol photons m$^{-2}$ s$^{-1}$ was evaluated by polynomial contrast (PC); the significance of PPFD levels at 2 % [$O_2$] was evaluated in terms of linear and quadratic polynomial orthogonal contrasts (POC) for unequal PPFD increments.

| Plant type | Tmt | [$O_2$] | PPFD | $\Delta^{18}O$ | $\Delta_{ca}$ | $\Delta_{ea}$ | $\delta_t$ | $\delta_e$ | $\theta$ |
|---|---|---|---|---|---|---|---|---|---|
| | | (mmol mol$^{-1}$) | ($\mu$mol photons m$^{-2}$ s$^{-1}$) | (‰) | (‰) | (‰) | (‰) | (‰) | |
| CA-18 | 1 | 200 | 1500 | 69.± 3 | 46 ± 8 | 49 ± 1 | -14.6 ± 1.2 | -3.1 ± 0.7 | 0.9 ± 0.1 |
| | 2 | 20 | 1500 | 43 ± 1 | 50 ± 12 | 52 ± 3 | -15.6 ± 3.4 | -2.3 ± 2.7 | 1.0 ± 0.2 |
| | 3 | 20 | 500 | 59 ± 5 | 69 ± 9 | 53 ± 1 | -14.0 ± 3.1 | -1.2 ± 0.8 | 1.3 ± 0.2 |
| | 4 | 20 | 200 | 91 ± 11 | 107 ± 13 | 51 ± 4 | -18.8 ± 8.9 | -3.5 ± 3.8 | 1.9 ± 0.3 |
| CA-39 | 1 | 200 | 1500 | 73 ± 4 | 49 ± 5 | 49 ± 1 | -13.4 ± 1.8 | -2.8 ± 1.0 | 1.0 ± 0.1 |
| | 2 | 20 | 1500 | 42 ± 3 | 49 ± 7 | 53 ± 1 | -12.7 ± 2.1 | -1.2 ± 1.2 | 0.9 ± 0.1 |
| | 3 | 20 | 500 | 65 ± 5 | 60 ± 11 | 53 ± 1 | -10.9 ± 1.4 | -0.7 ± 0.9 | 1.1 ± 0.2 |
| | 4 | 20 | 200 | 119 ± 12 | 67 ± 13 | 54 ± 1 | -9.8 ± 2.1 | -0.6 ± 1.1 | 1.2 ± 0.2 |
| WT | 1 | 200 | 1500 | 74 ± 4 | 35 ± 2 | 49 ± 2 | -13.7 ± 2.7 | -2.8 ± 1.7 | 0.7 ± 0.0 |
| | 2 | 20 | 1500 | 39 ± 3 | 54 ± 9 | 54 ± 0 | -15.1 ± 1.7 | -0.7 ± 0.7 | 1.0 ± 0.2 |
| | 3 | 20 | 500 | 59 ± 10 | 58 ± 7 | 52 ± 3 | -13.6 ± 1.9 | 0.9 ± 1.2 | 1.1 ± 0.1 |
| | 4 | 20 | 200 | 102 ± 30 | 80 ± 9 | 54 ± 1 | -11.2 ± 2.5 | -0.2 ± 1.1 | 1.4 ± 0.1 |
| | *Significance* | | *Plant type* | P= 0.58 | P= 0.26 | P= 0.86 | P= 0.60 | P= 0.63 | P= 0.24 |
| | | | *Tmt* | P< 0.0001 | P< 0.0001 | P= 0.006 | P= 0.86 | P= 0.14 | P= 0.0004 |
| | | | *Plant type * Tmt* | P= 0.70 | P= 0.28 | P= 0.80 | P= 0.63 | P= 0.90 | P= 0.41 |
| | | *Tmts 1-2* | *PC* | P< 0.0001 | P= 0.01 | P= 0.82 | P= 0.83 | P= 0.30 | P= 0.02 |
| | | *Tmts 2-3-4* | *POC linear* | P= 0.74 | P= 0.02 | P= 0.03 | P= 0.49 | P= 0.03 | P= 0.05 |
| | | | *POC quadratic* | P= 0.0001 | P= 0.64 | P= 0.01 | P= 0.66 | P= 0.35 | P= 0.92 |
