## Supplemental File S4 for "A role for neutral variation in the evolution of C_4_ photosynthesis"

### Phylogenetic tree of CA orthogroup

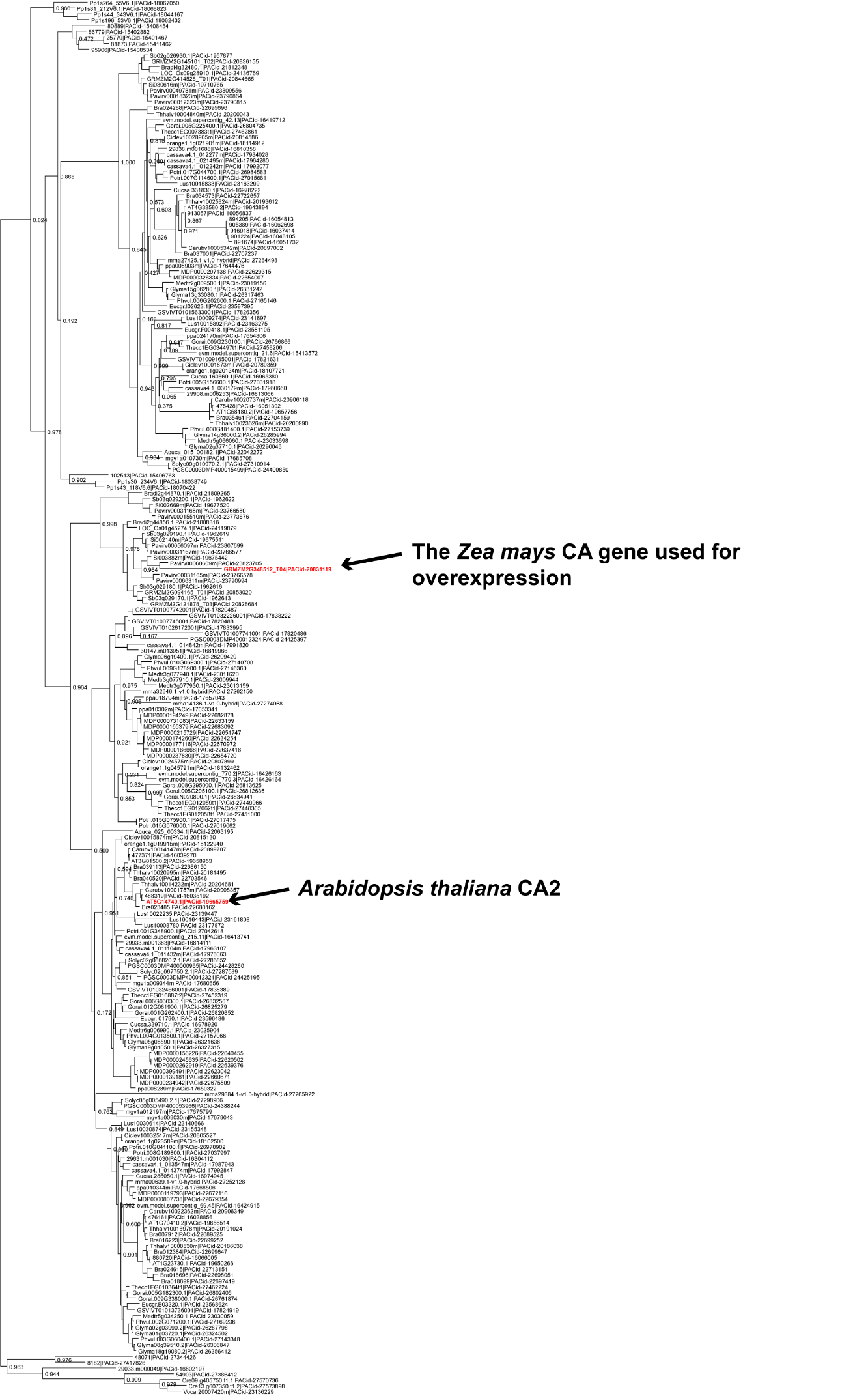

### Phylogenetic tree of PEPC orthogroup

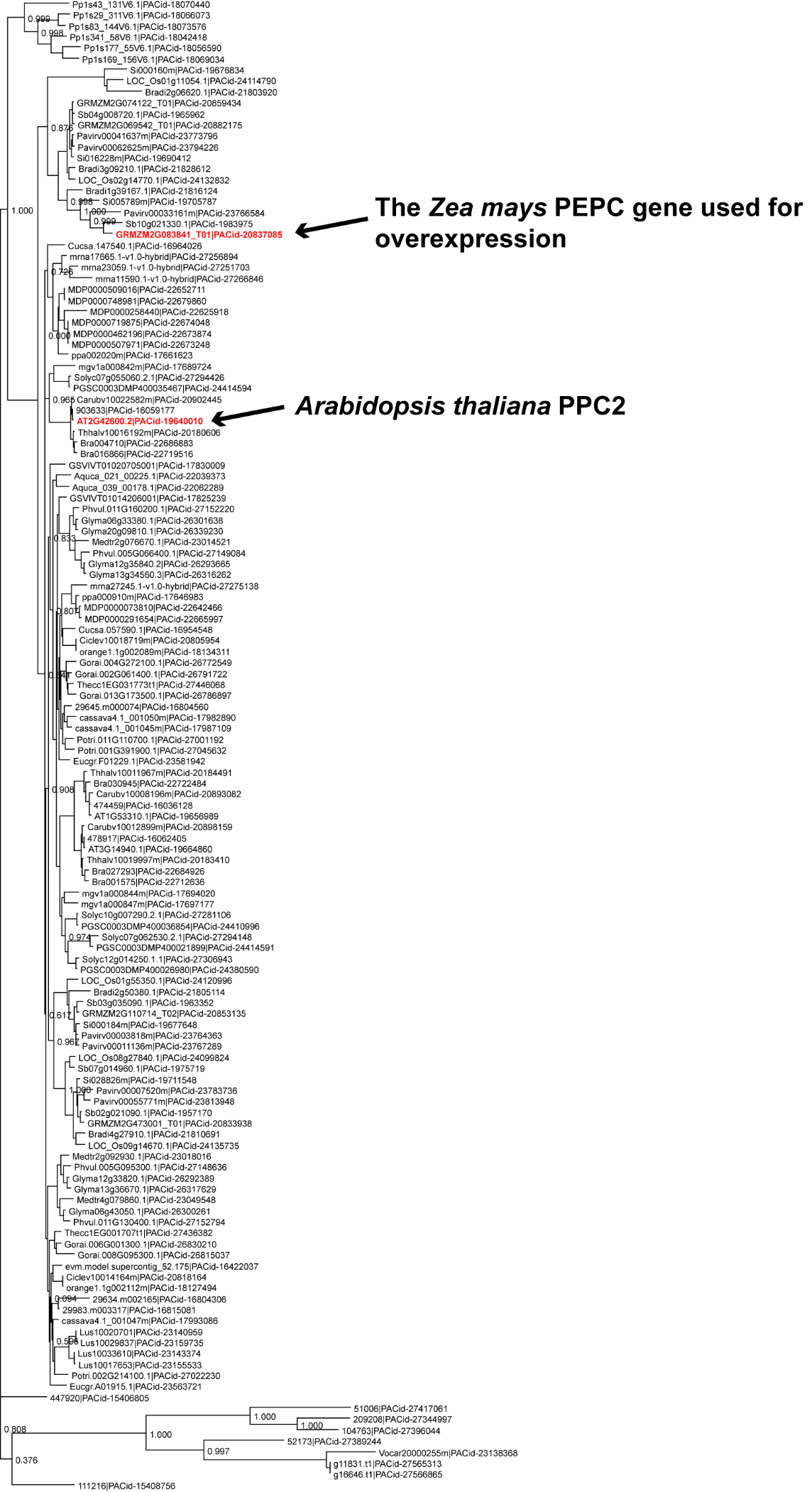

### Phylogenetic tree of MDH orthogroup

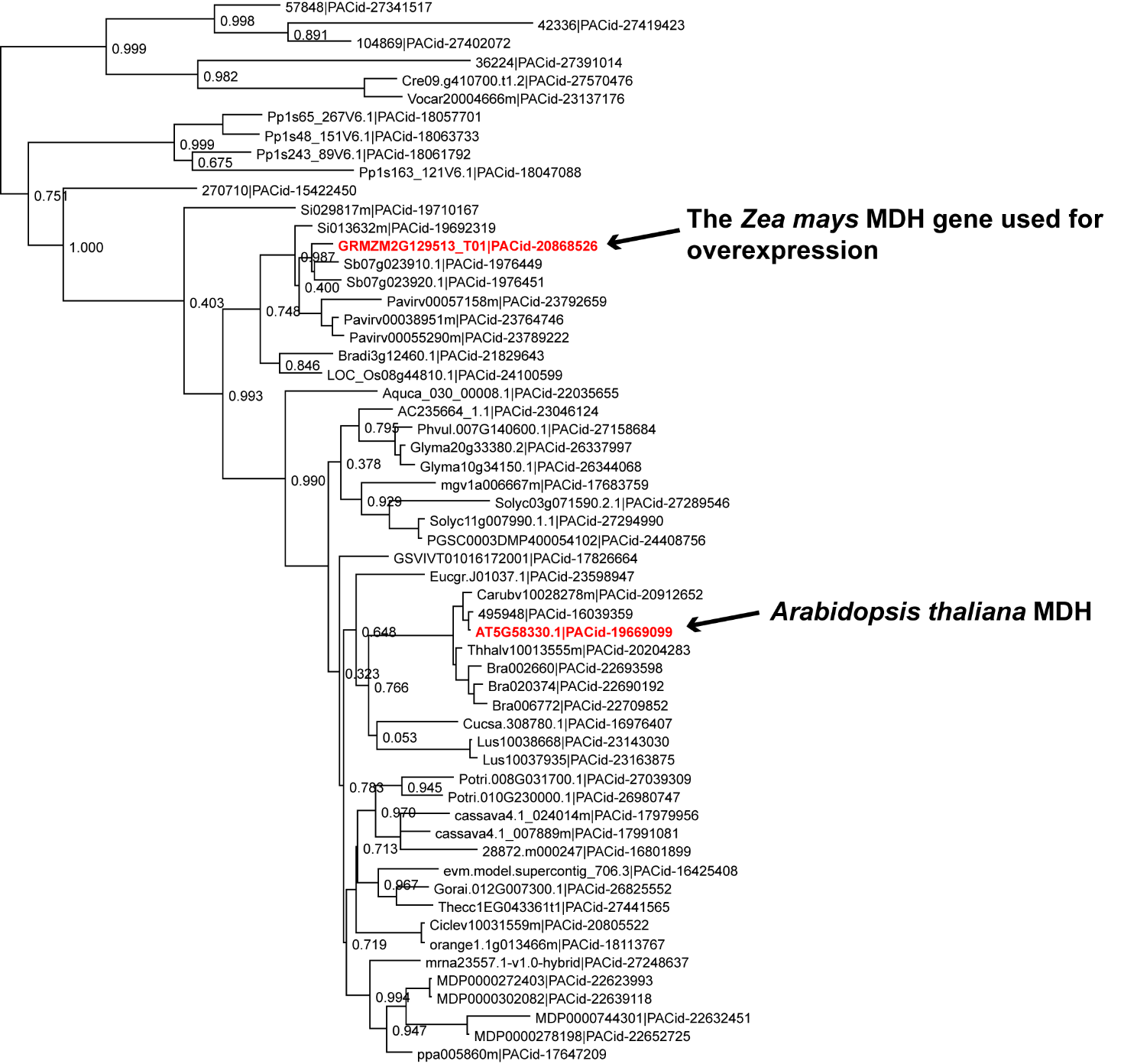

### Phylogenetic tree of NADP-ME orthogroup

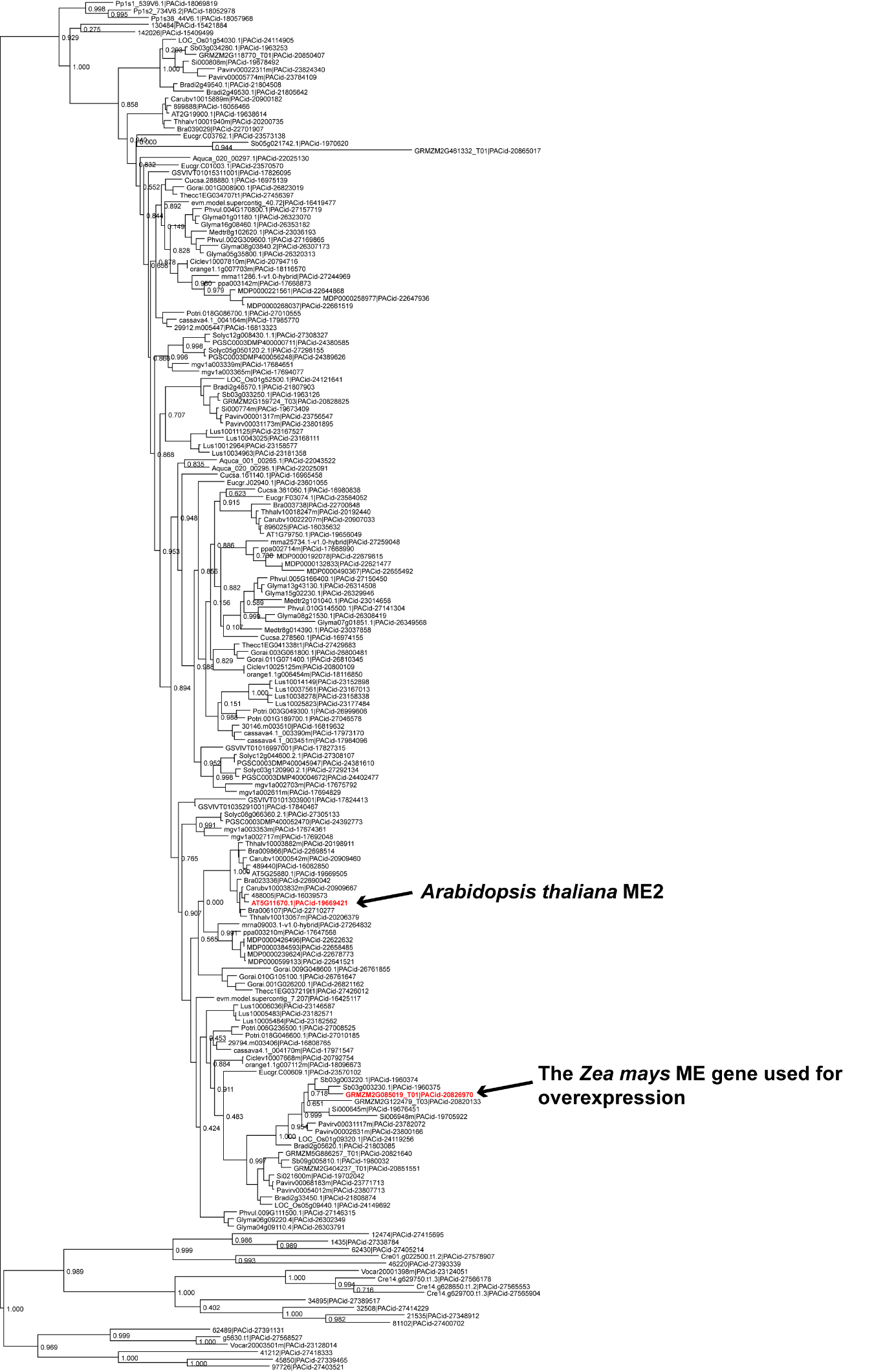

### Phylogenetic tree of PPDK orthogroup

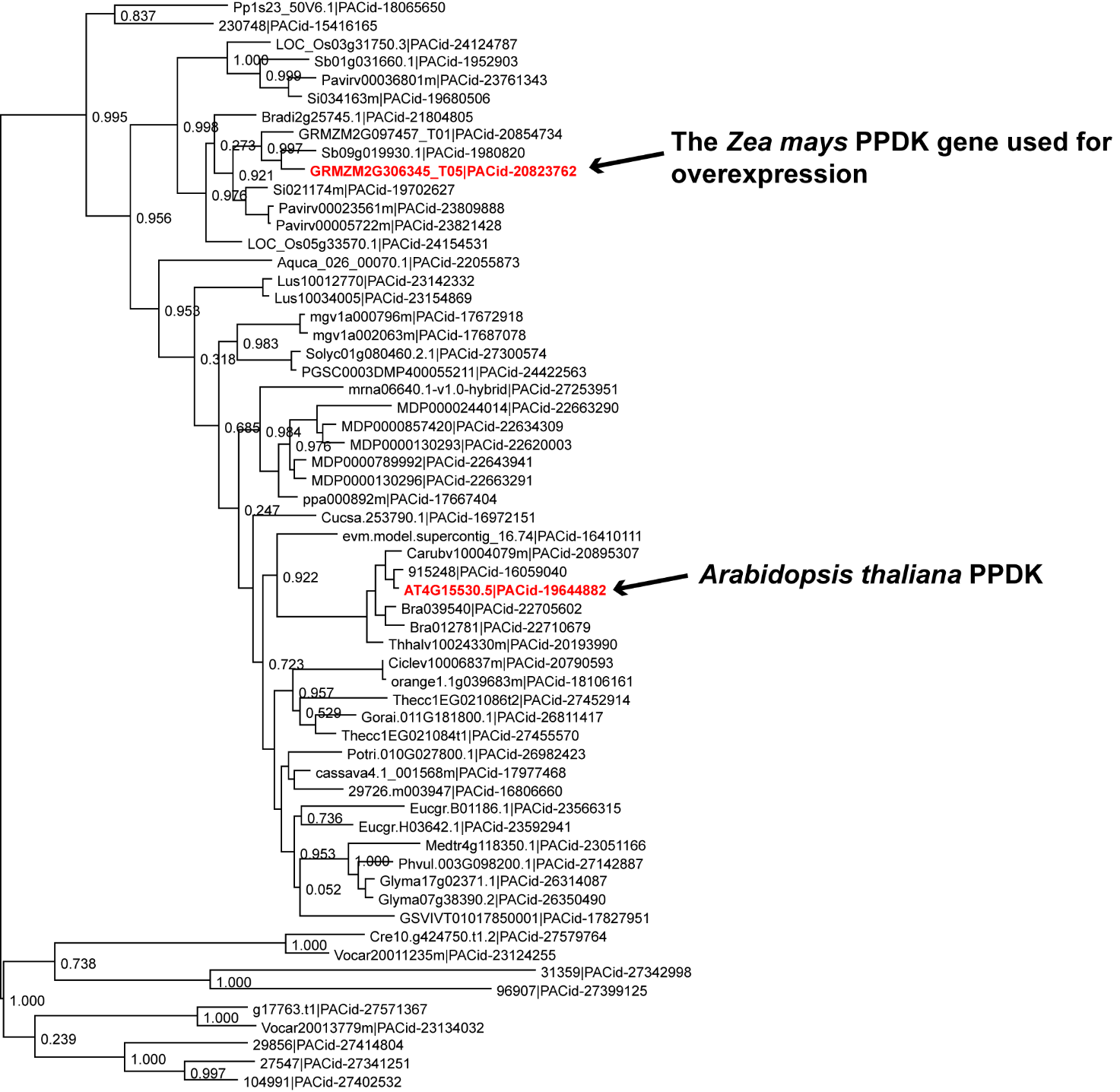
